# Large-scale single-cell mapping of concealed mitochondrial ageing

**DOI:** 10.64898/2026.07.14.738425

**Authors:** Hetvi Jethwani, Kamal Sukandar, Alistair P. Green, Ferdinando Insalata, Zoe Golder, Katherine Schon, Becca Asquith, Patrick F. Chinnery, Nick S. Jones

**Author notes:** These authors contributed equally (Hetvi Jethwani, Kamal Sukandar). These authors contributed equally (Alistair P. Green, Ferdinando Insalata).

## Abstract

Mitochondrial DNA (mtDNA) mutations have been linked to ageing. We find that the mtDNA mutations detected in routine bulk sequencing are greatly outnumbered by a class of concealed mutations that, despite being in a proliferative tissue, show little negative selection. Combining new scATAC-seq experiments with existing scRNA-seq, we map mtDNA mutations across 2.7 million cells from 311 individuals, covering 11 proliferative cell types and 5 tissues. We find that low-prevalence mutations (in ***≤***1% of assayed cells), concealed from routine bulk sequencing, accumulate markedly with age, with mutations unique to single cells (cryptic) predominating. We find a near universal accumulation rate across cell types and tissues, with, by age 50, approximately 40% of cells having at least one mutation affecting 30% of cellular mtDNA or more. At a correlative level we find that human age can be predicted from these concealed mutations but there is evidence of, possibly physiological, inter-individual variation. These low-prevalence mutations (unlike mutations tracking larger clones) show limited negative selection and coincide with gene-expression changes marking dysfunction.

## Main

Ageing is characterised by a set of interdependent molecular, cellular, and systemic hallmarks, including mitochondrial dysfunction [1]. Mitochondria are cellular power-stations and manufactories that contain a circular genome - mitochondrial DNA (mtDNA) - and play key roles in energy production, cell signalling, differentiation, and apoptosis pathways [2–5]. Over time, mitochondrial function declines, coinciding with the accumulation of somatic mtDNA mutations [6–9]. While inherited (germline) mtDNA mutations are already known to cause a large class of diseases [10–12], the functional relevance of somatic mtDNA mutations remains understudied [6–9]. Notably, mitochondrial mutator mouse models with an enhanced mtDNA mutation rate developed accelerated ageing phenotypes across tissues, such as the heart, muscle, brain, and colonic stem cells [13–16]. Specific to immune cells, mitochondrial dysfunction has been closely linked to senescent and exhausted T cells, triggering a cascade of age-associated dysfunctions [17–19]. However, whether somatic mtDNA mutations contribute to human ageing remains poorly understood.

At the cellular level, each cell has hundreds to thousands of mtDNA molecules [20, 21], allowing mtDNA mutations to exist in a state of heteroplasmy: the coexistence of wild-type and mutated mtDNA molecules within the same cell. At heteroplasmy of 60-90%, mtDNA mutations typically reach physiologically relevant levels that disrupt oxidative phosphorylation [22–24], although even lower heteroplasmy (∼30%) can have clear functional consequences [24, 25]. While it has been shown in bulk samples that both nuclear and mtDNA mutations increase with age, it is unclear whether mtDNA mutations reach physiologically high levels within individual proliferative cells during the human lifetime [26, 27].

A recent study on bulk sequencing of thousands of individuals has shown that negatively selected mtDNA mutations accumulate with age, likely within large clones, and are passive markers of clonal haematopoiesis with evidence that the mutations are likely not having a functional effect in their clones [28]. This study gives an important account of mtDNA mutation with age in large clones but it also exposes a key limitation of bulk approaches: they cannot identify mutations at high heteroplasmy which are found in small clones. Mutations contained in small fractions of the population are *concealed* from routine bulk approaches: it is the character of these concealed mutations that is the focus of our study. While the general view in the field (informed by bulk data and mitochondrial disease mutations) is that mutations in proliferative tissue are typically subject to purification [29, 30] how broadly this applies to concealed mutations in small clones is open.

We define the ‘prevalence’ of a mutant mtDNA site as the percentage of all sampled cells where the mutant site is detected (restricting to same-type cells where the site is covered, see Methods). We term mutations ‘low-prevalence’ if their prevalence is below or equal to 1% and ‘cryptic’ if the mutation is found in only one cell in a sample [26, 27]. We call these mutations Cryptic and Low-Prevalence mtDNA mutations (CLPs).

We present a large-scale single-cell analysis, in hundreds of donors, of rare somatic mutations, concealed from routine bulk sequencing, using new and published scRNA-seq and scATAC-seq data. We analyse data from 311 donors, comprising more than 2.7 million cells before filtering, across five tissues, two species, and 11 cell types. We show that CLPs are the most abundant class of mutations, comprising approximately 80-90% of the repertoire: almost all the mutations are cryptic. The heteroplasmy distribution of CLPs shows a marked shift to higher levels with age across proliferative cell types and shows little evidence of selection. By contrast, bulk-detectable prevalent mutations (at *>*1% of the sampled population) show weaker evidence of age accumulation and do show pronounced evidence of negative selection. CLP load per cell increases at similar rates across cell types, with 40% of cells harbouring a CLP by age 50. Levels of CLPs predict human age (reaching high heteroplasmy levels in late life) indicating relevance as a single-cell clock, and further we see marked inter-individual variation in inferred mitochondrial age. We see evidence that late-life individuals have a lower level of CLPs than is predicted from early-life accumulation rates: we exclude routes for selection within cells or within individuals. Despite CLPs showing limited evidence of selection their presence coincides with transcriptional markers of stress, quiescence and mitochondrial dysfunction in T cells.

## Results

### A large-scale inference pipeline shows cryptic and low prevalence mutations are abundant across proliferative cell types and tissues, and accumulate markedly with age, unlike prevalent mutations

In addition to our own multiomics combining ATAC and RNA-seq, we use 12 further scRNA-seq datasets to study the variation of cryptic and low-prevalence mtDNA mutations in human ageing (Supplementary Table 1). We define heteroplasmy (HF) of a given mtDNA mutation within a single cell as the fraction of mutated reads relative to the total number of aligned reads at the corresponding genome site. Heteroplasmy levels modulate the functional consequences of mtDNA mutations, with higher levels (e.g. HF≥30%) increasing the probability of cellular dysfunction [23–25]. We therefore restrict analysis to low-prevalence mutations with HF≥30% which also allows us to control for PCR-induced artefacts [31]. We thus only consider CLPs as those with HF≥30%. The age-related trends we report are unlikely to be technical errors, are robust to additive noise, and are independently validated using scATAC-seq (see Methods and Supplementary S1).

We find that CLPs are the most abundant class of distinct mutations in proliferative tissues, accounting for approximately 80-90% of the distinct mutation repertoire. For instance, for T cells obtained from 347 samples across 6 datasets, we see that ∼87% of the unique mutations are CLPs (Fig.1C for human T cells, and Supplementary S2.2 for other human cell types).

**Fig. 1.**
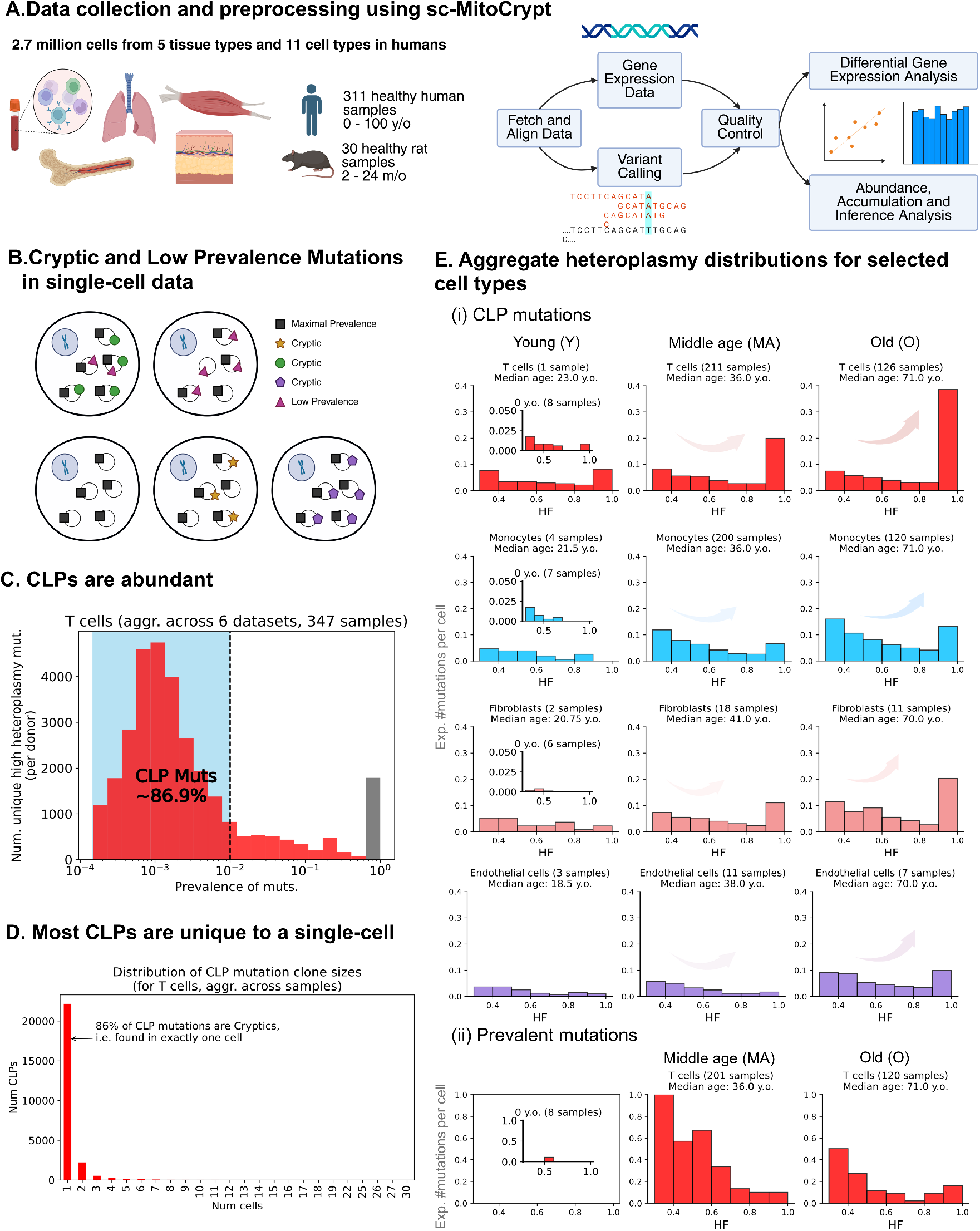
Large-scale single-cell RNA sequencing data analysis across 311 donors reveals the abundance of cryptic and low-prevalence mtDNA mutations (CLPs), uncovering universal dynamics indicative of shifts to higher heteroplasmies with age, in contrast to limited mutation accumulation among high prevalence variants: A. Overview of the sc-MitoCrypt pipeline, used to call variants from scRNA-seq and scATAC-seq data: 2.7 million cells (pre-filtering) from 311 donors across five tissues in humans, and 30 samples from rats; B. Diagram that distinguishes CLPs localised to a few cells from maximally prevalent mutations; C. CLPs are the most abundant class of distinct mutations in T cells, forming 80-90% of the unique mutations detected aggregated across 347 samples across six datasets. D. 86% of the unique CLPs are cryptic, i.e. unique to a single cell; E. Heteroplasmy distributions of CLPs in E(i) show a universal heteroplasmy shift with age across proliferative cell types, in contrast to haplotype-excluded prevalent mutations show in E(ii) that exhibit limited mutation accumulation with age. We show instances of the heteroplasmy distribution for T, monocytes, fibroblasts and endothelial cells for E(i) and T cells for E(ii). We normalise the heteroplasmy distribution by the total bases passing and extrapolate to the entire genome, to give an indicative scale, to get the expected number of mutations per cell in a given heteroplasmy bin (Methods). Extended figures for C, D and E are in Supplementary S2.

Most CLPs are unique to single cells (75−96%): for instance, in T cells across all datasets, ∼86% of CLPs are cryptic (Fig.1D for human T cells and Supplementary S2.2 for other human cell types). In bulk RNA-seq analyses, CLPs are typically filtered out as noise due to low pseudobulk heteroplasmy and hence, go largely undetected. The predominance of cryptic CLPs suggests that CLPs accumulate in distinct clones in the sample–consistent with the highly polyclonal character of many proliferative cell types [32]. We validate the polyclonality assumption by investigating the co-occurrence of TCR clonotypes with CLPs using scTCR-seq data in Supplementary S3.

Next, we investigate age-associated shifts in the mutational landscape of CLPs by examining the distribution of heteroplasmic and homoplasmic CLPs across four age groups: C (cord blood), Y (young), MA (middle age) and O (old). We calculate the aggregate heteroplasmy distribution by combining CLPs across donors for each cell type and age group pair (Methods). The aggregate heteroplasmy distribution of CLPs consistently shifts to higher heteroplasmies with age across cell types and species (Fig.1E(i) for human T cells, monocytes, fibroblasts, and endothelial cells, and Supplementary Discussion S2.3 for other human and rat cell types). A similar heteroplasmic shift is observed in post-mitotic tissues and is consonant with the corresponding coalescent theory [26, 27]. Despite the observed universal shift towards homoplasmy with age, we observe cell type-specific heterogeneity in the heteroplasmy distribution. For instance, T cell data can exhibit ∼40% homoplasmic CLPs in the MA group, in contrast to endothelial cell data that harbours relatively less homoplasmic CLPs in the same group, at about 8% (Fig.1E(i)).

We use the term ‘prevalent’ to refer to mtDNA mutations, with prevalence *>* 1% and *<* 90%, to help us visualize age-associated somatic accumulation. Mitochondrial mutations with ≥ 90% prevalence reflect either haplotype or germline heteroplasmies and are referred to as Maximally prevalent (MP). It is unclear whether prevalent mtDNA mutations show age-associated accumulation, although we expect the signal to be weak given that large clones are small in number per capita [33] (Fig.1E(ii) for human T cells, and Supplementary S2.3 for other human cell types). Our sampling of prevalent mutations is limited by clonal deduplication and low coverage of the mitochondrial genome in scRNA-seq data (Supplementary S1 and Methods).

### Levels of CLPs predict age, synchronously hit high levels across cell types in mid-life, and mark periods of high mortality

For each individual, we estimate a scaled frequency of observed CLPs per base to obtain an indicative statistic for the expected number of unique CLPs in a cell or CLP load (Methods). We observe an age-related increase in the CLP load across diverse cell types and tissues in humans (Fig.2A). An age-associated positive slope is observed for 9/11 cell types, namely T cells (slope = 0.00696, p = 4.8e-62), NK cells (0.00497, p = 1.133e-05), B cells (0.00655, p = 2.107e-06), monocytes (0.00724, p = 2.550e-27), macrophages (0.00611, p = 1.183e-04), fibroblasts (0.00781, p = 1.803e-06), endothelial cells (0.00502, p = 8.985e-03), MUSC (0.00578, p = 3.812e-02), and epithelial cells (0.00952, p = 1.392e-02). We have much less data for HSCs and keratinocytes, making our gradient estimate less certain (p = 0.074 and p = 0.056, respectively). Nonetheless, despite the small sample size, their CLP load is significantly shifted away from zero (one-sided t-test p: 0.0002 and p: 1.089e-5). Levels of CLPs predict age and are thus candidates for a single-cell ageing clock. Notably there are no parameters trained on chronological age and the statistic is derived from a physiologically tested and apparently cell-autonomous process. Although CLP loads follow the fitted age trend, there is an appreciable amount of variation around the regression line (Fig.2A). This is only partly reflected by limited sampling depth per individual in RNAseq data, and may also indicate genuine interindividual variation in CLP load, potentially linked to biology or environment, which we examine further in the next section and Supplementary S5.3.

**Fig. 2.**
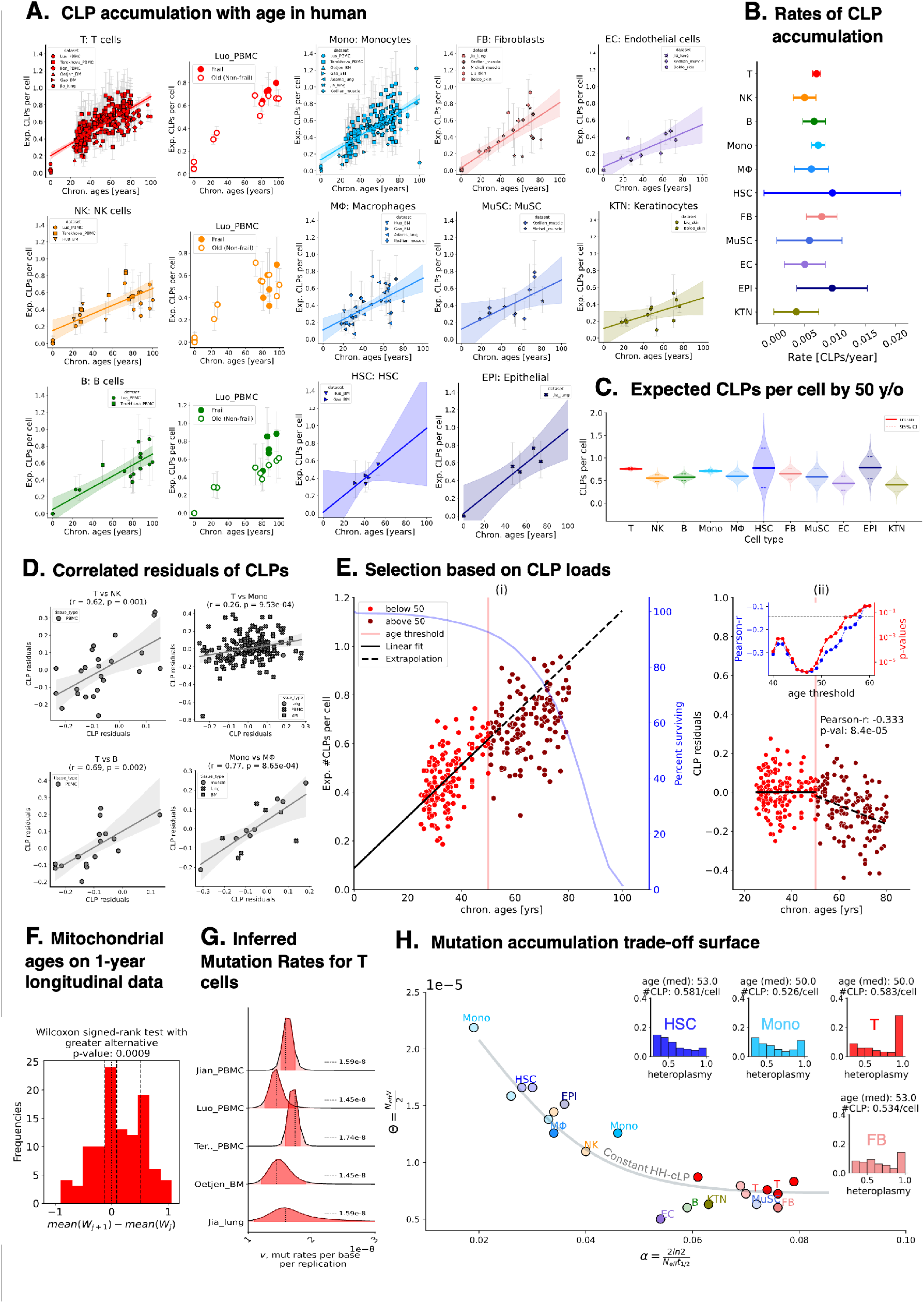
Universal accumulation of CLP across human cell and tissue types, with a late-life shift and a conserved absolute number per cell in late life. **A.** The inferred CLP loads, defined as the expected number of CLPs per cell, are shown as individual datapoints per donor across 11 human cell types (whiskers represent 95% CI). Specific to T, B and NK cells, we also show donors from Luo et al. [35] to compare CLP loads in frail and non-frail old individuals; **B.** Comparison of CLP accumulation rates (per year) across cell types, derived from linear fits using all available data points per cell type; **C.** Distributions of the expected number of CLPs by age 50, indicating comparable midlife CLP loads across cell types; **D.** We show strong positive correlations of CLP residuals between selected pairs of cell types from donors providing both cell types. CLP residuals were computed based on the linear fits shown in A; **E.** We present the evidence of selection using CLP loads in T cells (Terekhova et al. [34]): (i) Linear model fit using data from individuals below age 50 and extrapolated to those above 50, (ii) Residuals of CLP for individuals above 50 show a decreasing trend with age, in contrast to those below 50. The inset figure shows the robustness of this trend by sweeping the age thresholds. **F.**Distribution of differences in inferred mitochondrial ages *W* from samples collected from the same individual one year apart; **G.** Posterior distributions of mutation rates per base per replication *v* from model fitting to donors below 50 from five T cell datasets (3 PBMC, 1 BM, and 1 lung) lie in a consistent range, with the dotted lines representing the maximum a posteriori (MAP) estimate; **H.** Landscape of estimated ageing rates (*α*) and scaled mutation rates (Θ) across 21 cell-type–dataset pairs, revealing a strong trade-off between the two parameters. Estimates are primarily fit using donors aged ≤50 years to minimise selection effects (shown as fully opaque data points). Datasets with limited younger donors include all ages and are shown with 50% transparency. The inset figure shows the heteroplasmy distribution for selected cell types, demonstrating that despite differences in heteroplasmy distribution shape, a comparable number of CLPs is maintained by midlife. For example, T cells and fibroblasts—despite higher turnover rates than HSCs and monocytes—exhibit reduced scaled mutation rates Θ, resulting in similar CLP loads by midlife.

We find comparable rates of CLP accumulation across different proliferative cell types in humans: approximately 40% of cells harbour at least one CLP by age 50 (Fig.2B-C and Supplementary S2.1). A similar trend is observed in rats, but with substantially higher per-annum rates; by 24 months, ∼10% of rat cells harbour at least one CLP (Supplementary S2.7). The accumulation in a cell-type-independent manner is striking, given the marked variation in mtDNA copy number between cell types and the observed heterogeneity in heteroplasmy distribution dynamics across cell types– particularly between lymphoid and myeloid human lineages [20]. Next, we examine how CLP loads co-vary across cell types within individual donors. Donor-level CLP residuals are positively correlated between T and NK cells (r=0.62, p=0.01), T cells and monocytes (r=0.26, p=9.53e-4), and T and B cells (r=0.69, p=0.002) in PBMCs, consistent with a shared lineage (Fig.2D). We also observe strong covariation between monocytes and macrophages in human muscle, lung and bone marrows (Fig.2D). Thus, donors with higher-than-expected CLP loads in one blood cell type also tended to have higher loads in others, suggesting a coordinated accumulation of CLPs across cell types within each individual, that could be driven by donor-specific rates of accumulation.

We find evidence for a selection effect linking age and CLP load. We use a standard linear model on data from Terekhova et al. [34], for individuals below the age of 50, and extrapolate to find predicted CLP load values for individuals above 50. Individuals over 50 have lower CLP loads than predicted (Fig.2E(i)). Moreover, this deviation increases in a manner that is consistent with increasing selection with age (Fig.2E(ii)). The selection effect might be occurring at the cellular level or within individuals. Looking at both the dN/dS and the heteroplasmy distribution of CLPs, we do not see evidence that selection at a cellular scale is removing mutations in both younger and older individuals (Fig.3 and Supplementary S4.1). We further note that the heteroplasmy distribution of older individuals with low levels of CLPs resembles that of younger individuals with similar levels, indicating no distinctive process in these individuals (Supplementary S2.5).

**Fig. 3.**
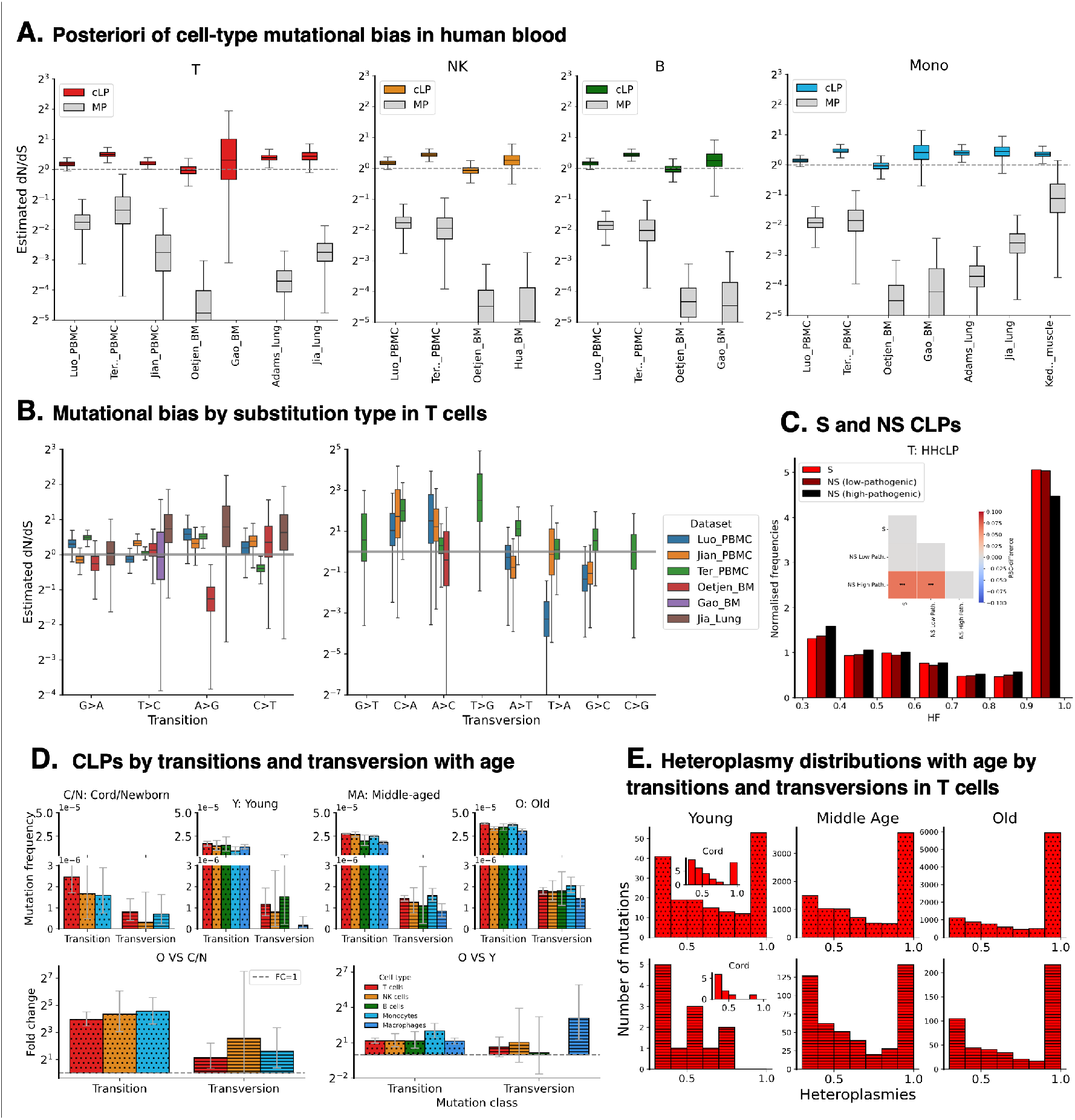
Limited evidence for selection in CLPs. **A.** Estimated dN/dS ratios across the different cell and tissue types, showing weak evidence of selection in CLPs, in contrast to maximally prevalent variants (in greys). For each dataset, cells of the same type were pooled across donors, and the number of synonymous (S) and non-synonymous (NS) variants was used to compute the posterior distribution of dN/dS. Posterior summaries are shown as median and interquartile range, with 95% credible intervals indicated by whiskers; **B.** In T cells, the procedure in (A) was repeated for each substitution type, as substitution types show different rates of occurrence that could bias the inference of overall dN/dS–even after this adjustment, still no evidence of selection was observed; **C.**Normalised heteroplasmy distribution of S, NS low pathogenic and high pathogenic mutations in the aggregated T cells across tissues and donors from five T cell datasets. We used MutPred to estimate the pathogenicity score. To compare the heteroplasmy distribution between S, low- and high-pathogenic NS, we show the computed RBC difference (**: p-values=1e-3); **D.**CLP load per base, i.e. mutation frequency, for transitions and transversions in four age groups (C: cord, 0 y.o., Y: young, 1-24 y.o., MA: middle-age, 24-60 y.o, and O: old, 60 y.o.). The colour represents the five cell types of blood as we exclude HSCs due to the donors having a narrow age range. The bottom panel shows the log2-fold change of mutation frequency between old and cord/newborn or young group, showing a significant increase in transversion CLPs in T, NK and monocyte at least when comparing between cord/newborn and old individuals; **E.** Heteroplasmy distribution of CLPs from 5 T cells datasets, disaggregated by transition (upper row) or transversion (lower row) in T cells, shows that both mutation types can segregate to higher heteroplasmies. RBC values are included to illustrate that while transversions reach higher heteroplasmies, their overall heteroplasmies remain less than those of transitions.

We present evidence that inter-individual variation in CLP load exists (next section). Eliminating obvious evidence for selection within individuals motivates future experiments exploring how this apparent late-life depletion might link to health or data-collection artefacts. A small amount of frailty data with 6 cases and 5 controls finds levels of CLPs in T cells and B cells (though not in NK cells) significantly higher in the frail individuals (T cells: Pearson corr=0.69, adj-p=2.5e-2; NK cells: Pearson corr=0.10, adj-p=0.77; B cells: Pearson corr=0.67, adj-p=3.0e-2). The idea of CLP load being relevant for individual health fits with the consistent control of their levels with age across cells and tissues observed in Fig.2B and C.

### Mutation accumulation is consonant with coalescent theory and dynamics conserve CLP load by mid-late life

The evidence we find of marked polyclonality (Fig.1C and Supplementary S3) is compatible with our knowledge of proliferative cell types [36, 37]. This allows us to use a model that assigns independent cell lineages to cryptic mutations [26, 27]. For each donor, we fit the model to the heteroplasmy distribution of CLP using Bayesian inference (Supplementary S5).

We can infer a mutation count per generation Θ which is broadly consistent across T-cell data from three different tissues, namely PBMCs [34, 35, 38], bone marrow [39], and lung [40]. Under the assumptions that the mtDNA copy number *N*_eff_ = 1000 and Θ = (*N*_eff_ · *v*)*/*2, we estimate the per-base per-replication mutation rate *v* to range from ≈ 1.45 × 10*^−^*^8^ to 1.76 × 10*^−^*^8^, in agreement with previous literature [26, 27, 41] (Fig.2G). We can also infer an effective mitochondrial age *W* of individuals in cross-sectional datasets, which increases in a clock-like manner with donor age at a rate of 0.051–0.069 units per year (Supplementary S5.3). Analysis of the one-year longitudinal data from Terekhova et al. [34] further shows that individuals gradually accumulate CLPs over time (Fig.2F). As in the earlier CLP-load analysis, we observe substantial variation around the fitted mitochondrial-age line with age (Fig.2A). Consistently, the posterior for the dispersion parameter of the ageing rate *α* is significantly above zero (Supplementary S5.3), indicating inter-individual variation in mitochondrial ages: whether this is linked to biological, environmental or unaccounted technical variation is open.

In Fig.1D and Supplementary S2.3, we observe that the data show different modes of accumulation of CLPs: some cell types show a rapid fixation of homoplasmies (e.g. T cells) while others show only a gradual increase to higher heteroplasmies with limited fixation of homoplasmies (e.g. Monocytes). Despite these different modes, we observe that levels of CLPs are similar by age 50 (Fig.2B,C) across cell types. In principle, this could be achieved by varying the effective copy number *N*_eff_, mutation rate per base per molecular replication *v*, or the turnover rate (closely linked to *t*_1*/*2_, which captures both cell division rate and mtDNA birth-date rate). However, cell division alone is unlikely to explain the age-related shift towards higher heteroplasmies relative to baseline mitochondrial birth and death 2 ln 2 (Supplementary S5.3). Plotting Θ versus 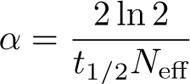 points to a common trade-off surface (Fig.2H). Assuming that the mutation rate per base per replication is constant across cell types, then the variation in Θ is driven by variation in *N*_eff_. Cells with high *N*_eff_ will limit the number of CLPs (since high *N*_eff_ can slow the spread of mutations to higher heteroplasmies) thereby limiting their possible physiological effects. The effective population size *N*_eff_ need not map onto the true number of mtDNA because of mitochondrial network fragmentation and subpopulation structure [42, 43].

### CLPs are broadly unselected while widespread mutations show strong negative selection

We assess selective pressure on variants in mitochondrial protein-coding genes by inferring the ratio of Non-Synonymous (NS) to Synonymous (S) mutation rates, defined as dN/dS ratio (Methods). As scRNA-seq data is underpowered for dN/dS inference at the cell or individual-level, we pool variants from cells of the same type across donors in a given dataset to assess the overall selective pressure of Non-Synonymous (NS) CLPs.

The dN/dS ratio of CLPs is largely consistent with neutrality. Across human proliferative tissues and cell types, we find that most datasets show no evidence of selection acting on NS CLPs (Fig.3A; e.g. T-cell Luo PBMC: median dN/dS=1.09; 95%CI 0.92-1.17, Jian PBMC: median dN/dS=1.12, 95% CI 0.97-1.20, Oetjen BM: median dN/dS: 0.91; 95% CI 0.62-1.2, Gao BM: median dN/dS=0.80, 95% CI 0.13-4, Jia lung: median dN/dS=1.30, 95% CI 0.90-2.25, and Adams lung: median dN/dS=1.31, 95% CI 0.97-1.85). However, only one dataset shows a slight elevation in the median of dN/dS, which suggests weak positive selection acting on NS CLPs (T-cell Terekhova PBMC: median dN/dS=1.45, 95% CI 1.06-1.80). By contrast, the dN/dS ratios of variants that are maximally prevalent (MP) are significantly below 1 (Fig.3A; e.g. T-cell Luo PBMC: median dN/dS=0.29, 95% CI 0.11-0.76, see Fig.3A for other cell types), consistent with purifying selection at the organismal level. Prevalent mutations also show evidence of negative selection, at least in the 1–10% prevalence range where we have sufficient data to estimate dN/dS (Supplementary Fig.S4B). Together, these results suggest a progression in which CLPs are largely neutral, whereas mutations that expand to higher prevalence are constrained by negative selection.

We compute the dN/dS separately for each of the 12 substitution types to account for the substantial variation in mutation frequencies by substitution type. We continue to find weak to no evidence of selection acting on CLPs across substitution type, with almost all of dN/dS posteriors including neutrality within their 95% CIs (Fig.3B for T cells), indicating that CLPs evolve neutrally in a manner robust to mutation-spectrum effects. When stratifying NS variants by predicted pathogenicity (Methods), highly pathogenic variants show a slight but statistically significant skew toward lower heteroplasmy levels (Fig.3C).

While dN/dS analyses indicate that CLPs in protein-coding regions are largely neutral, we next examine the full CLPs spectrum to assess how different substitution classes accumulate and segregate with age. Stratifying variants into transitions and transversions across major human blood cell types, we find that transitions account for ∼90–95% of CLPs and that their load increases with age (Fig.3D; T cells: median log2-FC=4.0, 95% CI [3.5,4.5]; NK cells: median log2-FC=4.3, 95% CI [3.0,6.0]; and monocytes: median log2-FC=4.6, 95% CI [3.8,5.7]). Transversions are comparatively rare but show some evidence of age-related accumulation, although limited counts of transversions result in larger error bars compared to transitions. This trend is consistent with previous bulk studies using Duplex-seq on rare somatic mutations in mice [44–46], which have suggested that transversions are rare, but here we see evidence of accumulation with age (Fig.3D; T cells: median log2-FC=1.1, 95% CI [0.3,2.2]; NK cells: median log2-FC=2.5, 95% CI [0.1,6.7]; and monocytes: median log2-FC=1.6, 95% CI [0.3,3.4]). Our analysis of single-cell data, however, allows us to add another layer to the debate. Analysis of single-cell heteroplasmy distributions in T cells shows that transversions can also reach higher heteroplasmy levels with age, similar to transitions (Fig. 3E). Nonetheless, transversion heteroplasmy distributions remain significantly lower than those of transitions across all age groups (RBC-difference between transition vs transversions in young=0.06, Mann-Whitney U p-value=0.73; middle-age=0.14, Mann-Whitney U p-value=1.2e-5; old=0.19, Mann-Whitney U p-value=4e-7), indicating that the rate of their expansion might be constrained relative to transitions. This points to a model where there is heavy selection against the introduction of transversions into the coding genome as mentioned in Cote-L’Heureux et al. [46], but once introduced, they segregate relatively neutrally like other mutations.

### CLPs are associated with gene expression changes indicative of ageing and senescence in T cells

We perform differential gene expression analyses (DGE) to quantify the degree to which the presence of Non-Synonymous CLPs co-varies with gene expression changes indicative of ageing and senescence, using T cells from three PBMC datasets having an adequate sample size [34, 35, 38]. Although data from Luo et al. [35] and Terekhova et al. [34] contain a large number of cells in total, relatively few cells have good mitochondrial genome coverage limiting the number of cells we observe with CLP mutations and so reducing statistical power. For each dataset, we compare the set of cells with only Non-Synonymous CLPs with the set of cells without CLPs (Methods and Supplementary S6). We aggregate cells across donors, balancing age, sequencing depth and mitochondrial coverage. Our analysis reveals a gene expression signature coinciding with the presence of non-synonymous Cryptic and Low Prevalence mutations, including genes previously linked to ageing and senescence pathways. The aggregated analysis results in 45, 14 and 5 recurrent, significant differentially-expressed genes with an absolute log2 fold-change higher than 0.1 for data from Jian et al., Terekhova et al., and Luo et al. respectively [34, 35, 38]. In Figure 4, we plot recurrent significant genes from Jian, and highlight selected genes from Luo and Terekhova (Fig.4A), full list S6. Note that since we largely use 10x data, we do not observe all mutations in a cell, weakening the contrast between cases and controls.

**Fig. 4.**
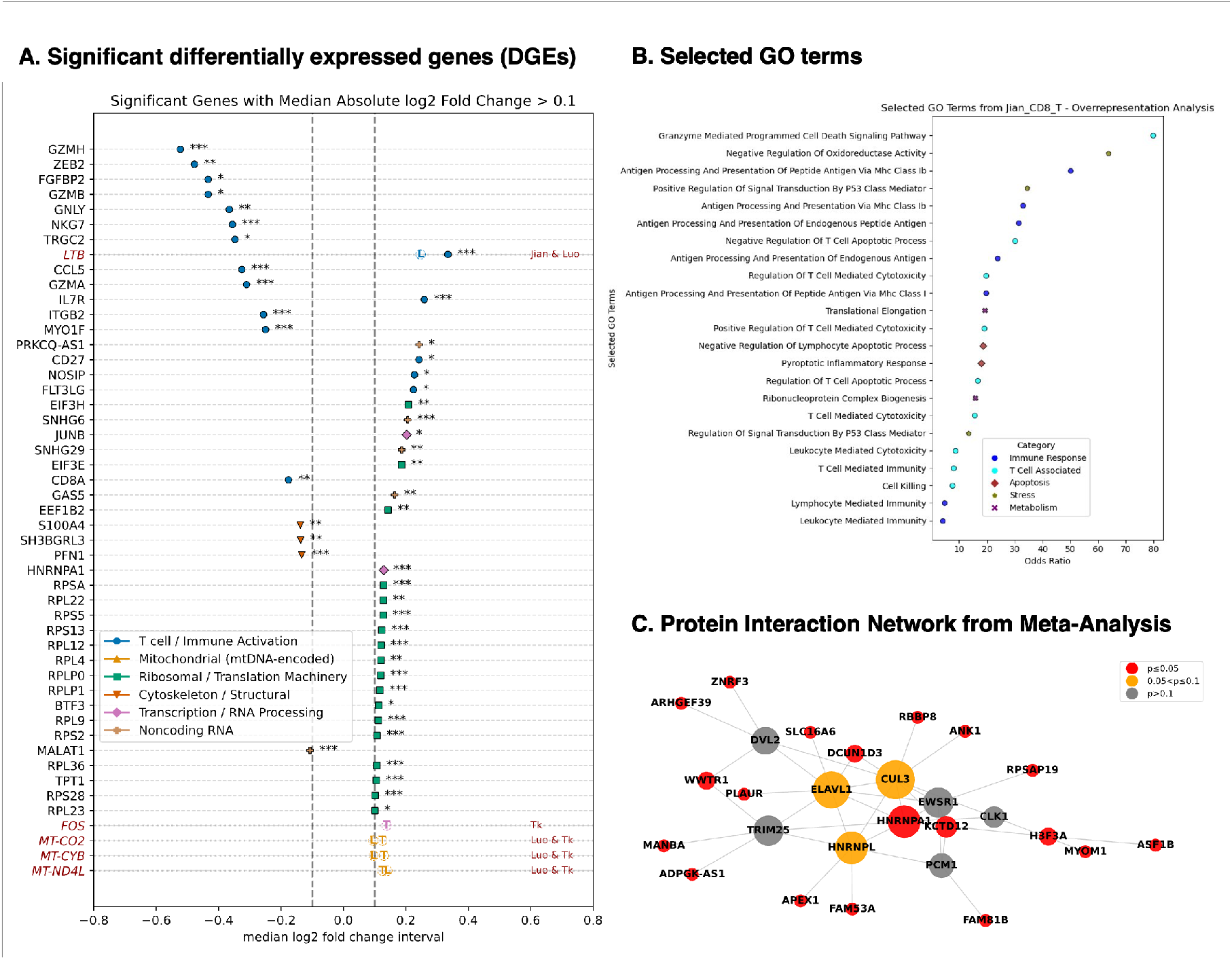
Transcriptional markers of stress and ageing covary with the presence of non-synonymous CLPs: **A.** Plot shows significant genes recurring in more than 30% of subsampling runs from T cells from Lu et al. [38]. For each gene, we plot the median log2 fold-change across subsampling runs. Each gene is coloured and marked by the type, the asterisks indicate strength of significance, representing the median adjusted p-values (Mann Whitney U test with importance subsampling, Benjamini-Hochberg correction; statistical significance denoted as * *p <* 0.05, ** *p <* 0.01, *** *p <* 0.0001). Genes with adjusted p value lower than 0.05 per sub-sampling run recurring in 30% or more subsampling runs, and median absolute log2 fold-changes above 0.1 have been plotted; Additionally we highlight the following genes recurring in significance across other datasets with the same analyses— MT-CO2, MT-CYB and MT-ND4L (significant in [34, 35]), LTB (significant in [35]), FOS (significant in [34]), see **Sec.**5 for details); **B.** Plot shows selected GO terms after performing a statistical overrepresentation test (Fisher’s exact test with FDR correction); Selected significant GO terms overrepresented in the gene list from [38] have been shown with the corresponding odds ratios, and the GO terms have been colored to indicate functional associations; **C.** Plot shows a protein-protein interaction network visualization of genes ranked by aggregate p-values obtained after performing a meta-analysis on individual-level Differential Gene Expression tests to increase statistical power. The network is obtained from SC-PPIN, and reveals ELAVL1, CUL3 and HNRNPL act as interaction hubs across proteins encoded by genes significant from the meta-analysis.

The aggregated analysis is consonant with altered T cell differentiation of cells harboring Non-Synonymous CLPs towards a quiescent/memory-like state. Several key genes associated with cytotoxic effector function, including NKG7, CCL5, CD8A, GZMB, GZMH, and GNLY, are significantly downregulated in CLP-mutant T cells; this may indicate impaired or reduced cytotoxic/effector T cell activity, or an imbalance in the cell sub-type composition. Note that IL7R and LTB are upregulated. IL7R is a hallmark of naive and central memory T cells and plays a key role in T cell survival and longevity. LTB, a gene that encodes a part of the Tumor Necrosis Factor cytokine superfamily of proteins and forms protein complexes present on the surface of activated CD4+ and CD8+ T cells, is upregulated across 2 datasets [47–49]. Increased LTB expression may indicate chronic immune activation or altered cytokine signaling. ZEB2 is significantly down-regulated, consistent with reduced terminal effector differentiation of CD8+ T cells an effect associated with impaired acute immune responses [50]. EEF1G, a translation elongation subunit, is upregulated; dysregulation of EEF1G and related components has been linked to age-associated translational remodeling and proteostatic disruption [51, 52]. FOS and JUNB, AP-1 components implicated in age-associated chromatin remodelling and identified as conserved inflammaging markers [53, 54], are upregulated, suggesting links between the presence of CLPs and transcriptional shifts that have been previously implicated in immune ageing.

Mitochondrial genes that are involved in OXPHOS-related pathways such as MT-ND4 and MT-CO3 are observed to be upregulated, consistent with previous observations that OXPHOS-related upregulation can be triggered as compensatory response to defects in energy production [55]. Additionally, short and long ribosomal protein-encoding genes such as RPL36A, RPS29, RPL36, and RPL39 are upregulated linking the presence of mtDNA mutations to translational shifts. Recent studies have also linked ribosomal dysfunction and ribosomal stress to cell cycle arrest, apoptosis and p53 signalling pathways [56, 57].

To identify biological processes associated with Non-Synonymous CLPs, we performed GO over-representation analysis using PANTHER on significantly differentially expressed genes from Lu [58] (Fisher’s exact test with FDR correction, Methods) [59]. The gene sets are enriched for GO biological pathways indicative of stress responses, metabolic processes, and immune response (Fig.4B). We present full gene lists and significant GO terms in Supplementary S6. We further perform Differential Gene Expression tests per donor to increase statistical power and gain further clarity on the observed functional effects. We derive *p*-values across cells with and without Non-Synonymous CLPs per individual, combine *p*-values using a meta-analysis procedure, and apply multiple hypothesis correction (see Methods). To visualize the individual meta-analysis, we construct a protein-protein interaction network over the resulting gene list using SC-PPIN (Fig.4). HNRNPA1, CUL3, HNRNPL, ELAVL1 (HuR) and TRIM25 emerge as central hubs-these genes play key roles in stress response mechanisms to maintain cellular homeostasis [60–65]. Interestingly, HNRNPA1 is important for the stability of Regulatory T cells, and defects in HNRNPA1 have been linked to age-associated diseases like ALS [66–68].

Our analysis links the presence of Non-Synonymous CLPs with genes and pathways associated with ageing, quiescence, and stress. We observe a shift towards more quiescent phenotypes in the aggregated DGEs. At the finer meta-analysis resolution, the hub-genes play a role in regulatory programs that may help mutated cells cope with stress. It is possible that T cells with a higher mutation load may preferentially adopt stress-tolerant expression profiles that favour survival and persistence.

### Results are calibrated across sequencing types and are biologically replicable

We ensure the calibration of our scRNA-seq findings by generating single-cell multi-omic ATAC data and employ mtscATAC-seq to achieve higher breadth of mtDNA coverage. We analyse 24 human peripheral blood samples obtained from 10 donors (age range 23–60 years), with 2–3 biological replicates available for 8 donors. Each sample contains predominantly T cells, with a median of 330 cells (Supplementary S7.1). Mitochondrial coverage is substantially higher from mtscATAC samples compared to scRNAseq-based datasets: in our data, a median of 95% of the mitochondrial genome is covered, with a median read depth of 20 per position (Fig.5A). Given almost full coverage, we observe a high variability in mtDNA mutational burden across different functional genome categories, including protein-coding genes, tRNA, rRNA, and non-coding regions. Ribosomal RNA regions have significantly higher levels of CLPs per base compared with the other three genomic categories (Fig.5A). Although much of the existing literature suggests that non-coding regions are expected to harbour higher mutation rates due to their hypermutability, recent single-cell studies have reported higher mutation burdens in rRNA than in the D-loop in certain tissues, including human blood [69]. We note that slightly lower mutational burdens in the non-coding region may be influenced by technical factors, such as the linearization of the circular mitochondrial genome during alignment, which can result in partial read loss at split regions.

**Fig. 5.**
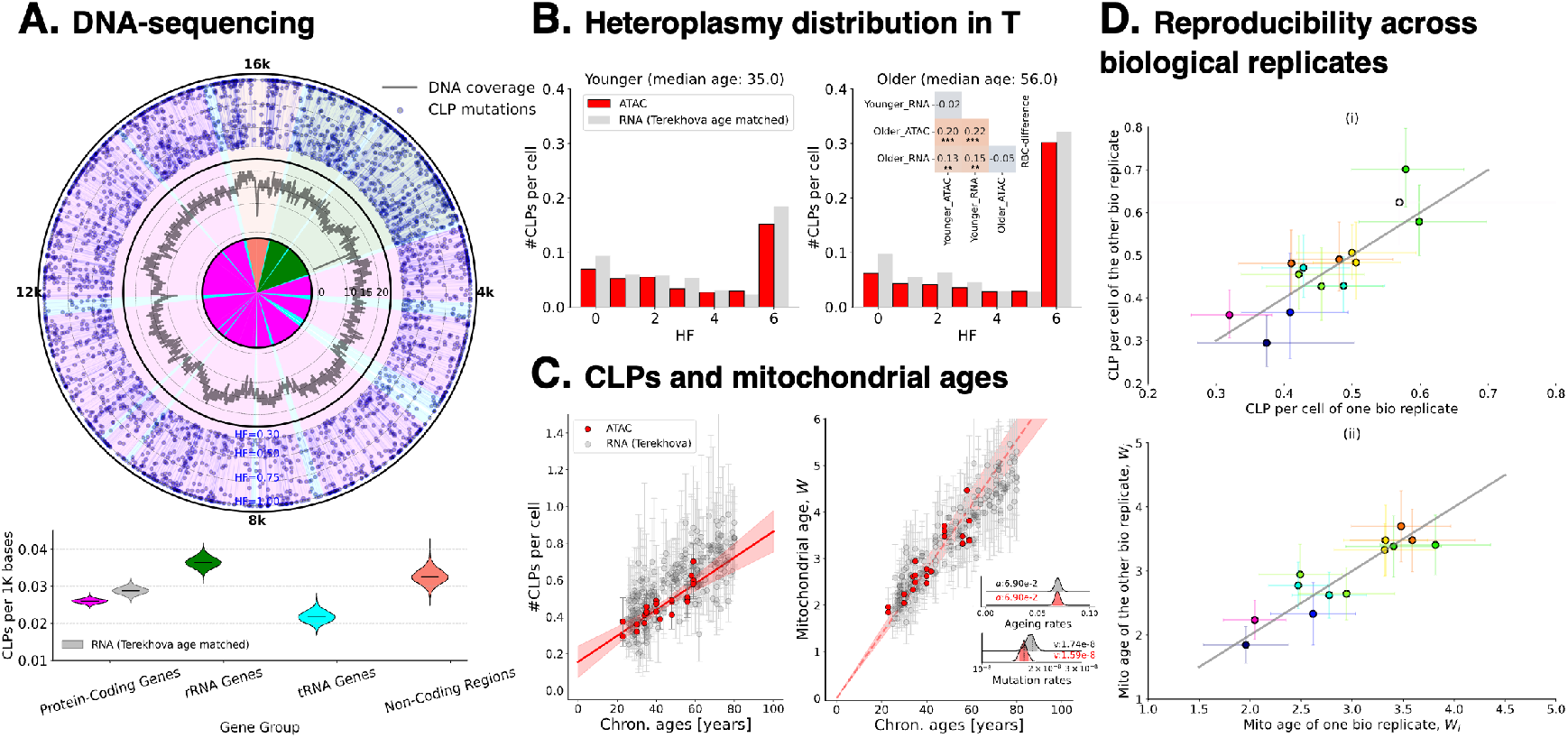
ATAC-seq data with higher mitochondrial genome coverage corroborate RNA-seq–derived results. **A.** Median of mitochondrial genome coverage with corresponding CLPs across 24 samples from 10 donors; colours denote mitochondrial gene groups. We quantify the amount of CLP load per 1k base across four gene groups, showing that ribosomal RNA exhibited a higher mutation load than the other groups; **B.** The heteroplasmy distribution of cells from younger (30–45 years, median 35) and older (45–60 years, median 56) individuals; **C.** Estimated CLP load per cell and inferred mitochondrial ages per individual, showing rates of increase comparable to RNA-seq–derived results (e.g. RNAseq from Terekhova, et al. in greys); **D.** Scatter plots of CLP loads and inferred mitochondrial ages from samples obtained from the same donors demonstrate high reproducibility across biological replicates.

We check the calibration of RNA-derived findings using the orthogonal modality and apply the same analytical framework to mtscATAC-seq data for T cells. Our results indicate a strong concor-dance between DNA- and RNA-based measurements of mitochondrial mutational dynamics in T-cell ageing. CLPs are similarly predominated by cryptic mutations (95%) and are largely neutral (Supplementary S7.1. Transitions are also a predominant form, but both transitions and transversions are segregated to higher heteroplasmies with age (Supplementary S7.1), consistent with RNAseq findings (Fig.3E). CLP heteroplasmy distributions differ significantly between young and old groups (Mann–Whitney U, p-value *<* 1 × 10*^−^*^22^; see Fig.5B in reds), indicating a shift towards higher heteroplasmies with age that is consistent with what observed via scRNA-seq in both magnitude and direction (see Fig.5B in grey). Next, we observe an age-dependent increase in the expected CLPs per cell, and inferred mitochondrial age for both sequencing modalities with comparable rates of accumulation (see Fig.5C). As a final check, we find that CLP load and inferred mitochondrial age are consistent across independent biological replicates, indicating that these metrics are robust to sampling variability and provide reliable measurements of mitochondrial mutational burden and age in ATAC-seq data (Fig.5D).

## Discussion

Studies of small numbers of individuals, cell types or in bulk have suggested the accumulation of mtDNA mutations with age [28, 41, 70–74]. Emerging evidence suggests that levels of mtDNA mutation do accumulate with age within large clones, and are detectable in bulk, but these mutations are negatively selected and are likely passive markers of clonal haematopoiesis [28]. Bulk assays conflate two distinct quantities: the prevalence of a mutation across a set of cells and its heteroplasmy in any particular cell. This distinction is central as a mutation might reach high heteroplasmy within a small set of cells while being concealed from routine bulk sequencing. By jointly resolving prevalence and heteroplasmy our study shows that this concealed, low prevalence, regime is where much of the age-associated mtDNA change in proliferative cells is found.

Using single-cell data, across 311 individuals and 11 cell types, and a new sc-MitoCrypt pipeline, we find that Cryptic and Low-Prevalence mtDNA mutations (CLPs, mutations in ≤ 1% of cells at heteroplasmy ≥ 30%) are the predominant form of mutation and that these mutations show a pronounced shift to higher heteroplasmy with age; by contrast more prevalent mutations are a minority of mutations and have a less clear pattern of accumulation with age (Fig.1C,D). An indicative value across assayed cell types is that 40% of cells have one or more CLPs by age 50 (Fig.2C); and we infer that 40% of cells have at least one CLP at homoplasmy by age 80 (Supplementary S5.3). We find that most CLPs are transitions consistent with a replication-error dominated process as a primary source of somatic mtDNA mutations [45]. We generate new mtscATAC-seq data, including biological replicates, to ensure calibration of our findings, and recapitulate the same patterns of CLP abundance and age-dependent accumulation (Fig.5).

We see an apparent convergence, across cell types and tissues, in the mid/late-life number of cryptic mtDNA mutations found at ≥30% heteroplasmy (Fig.1D, Fig.2A,B,C, Fig.5B,C). The convergence is consistent with a shared constraint on mutation tolerance for high heteroplasmy mtDNA mutations although the mechanisms of that intolerance remain uncharacterized. Age can be predicted from CLP levels, but we observe inter-individual variation which might be attributable to an unaccounted source of technical, genetic, environmental, or lifestyle variation (Supplementary S5.3).

We observe an apparent depletion of individuals with high levels of CLPs in late-life compared to early-life extrapolations and a small-study frailty effect which motivate future studies on longitudinal health effects of CLPs (Fig.2A,E).

High heteroplasmy mtDNA mutations can impair cell function [22]; with mutations observed to inhibit proliferation in culture, naive T-cells and mtDNA-mutant cybrids [75–78]. Our single-cell work on function is correlative, finding that in T cells, elevated mtDNA mutation burden might coincide with transcriptional markers of immunosenescence, translational remodelling and mitochondrial respiratory shifts linked to pro-inflammatory T-cell ageing (Fig.4A; [79]). We find limited evidence for selection amongst Non-Synonymous CLPs, unlike maximally prevalent Non-Synonymous mtDNA mutations which we find are strongly selected against (consonant with recent work [28]). This raises the testable possibility that age-associated CLPs create a purification burden linked to cellular expansion: a form of clonal blunting that complements the more familiar clonal expansions with ageing.

Our mapping of concealed Cryptic and Low-Prevalence mtDNA mutations establishes their widespread and consistently age-accumulating character across technologies, species, individuals longitudinally and in biological replicates, in multiple tissues and proliferative cell types. Vitally, we see that levels become appreciable by late life, paving the way for direct tests of their functional effects with age.

## Methods

We constructed a pipeline to align raw scRNA-seq data to the given reference genome for data generated by 10x Genomics Chromium Single Cell 3’ or 5’ Assays. The pipeline outputs a list of mtDNA variants and the gene expression matrix for each sample, which are subsequently used to infer the mutation state of individual cells. These outputs allow us to investigate the mtDNA mutational landscape and differential gene expression analysis in a cell-type-specific manner across our datasets. The pipeline is implemented using a combination of shell and Python scripts and is designed to run in parallel over a PBS-based high-performance computing cluster for computational efficiency.

### Data Collection

#### scRNA-seq sequencing data collection

We downloaded single-cell RNA sequencing data in .fastq files from publicly available repositories, including the Gene Expression Omnibus (GEO, https://www.ncbi.nlm.nih.gov/geo/), Synapse (https://www.synapse.org/Home:x), BioStudies (https://www.ebi.ac.uk/biostudies/), and Genome Sequence Archive (GSA, https://ngdc.cncb.ac.cn/gsa-human/). For raw data in GEO, we used fastq-dump command from SRA-toolkit to fetch metadata and to download raw sequencing files of samples or individuals of interest. For raw data in Synapse, we use the synapseclient python package to download raw sequencing files. For datasets stored in GSA, we used direct links provided on the website to copy the .fastq files to a local drive. A complete list of all accession IDs used in this manuscript is provided in Supplementary Table 1.

### Isolation of Peripheral Blood Mononuclear Cells for mtscATAC-seq

#### Cell isolation

Peripheral venous blood was collected into EDTA-coated tubes and processed within 30 mins of collection. Whole blood was diluted 1:1 with phosphate-buffered saline (PBS) containing 2% fetal bovine serum (FBS) and layered onto 15 mL of Lymphoprep (STEMCELL TECHNOLOGIES 18061) in SepMate-50 tubes (STEMCELL TECHNOLOGIES 85450). After centrifugation the upper layer containing PBMCs was poured into a fresh 50 mL conical tube and washed twice with PBS containing 2% FBS before the cells were counted, aliquoted, cryopreserved, and stored in liquid nitrogen until use.

#### Cell staining

Batches of cells were thawed, washed, and counted before being resuspended in staining buffer (Biolegend 420201) and blocked with Human TruStain FcX Blocking reagent (BioLegend 422301). Subsequently cells were incubated with TotalSeqA conjugated antibodies (Biolegend A0251, A0252, A0253) for 30 minutes. After washing and resuspension cells were stained with DAPI (Thermo D21490) and DRAQ5 (Thermo 65-0880-96) and live cells were sorted on an Influx Cell Sorter (BD Biosciences).

#### Cell fixation and permeabilization

Equal numbers of each sample were combined and prepared for 10X Genomics multiome ATAC and gene expression sequencing using a modified version of the 10X Genomics nuclei isolation protocol [80]. After fixing with formaldehyde (0.1% final concentration) cells were quenched with glycine (0.125 M final concentration) and subsequently washed. The pellet was resuspended and permeabilized on ice with 10 mM Tris-HCl pH 7.4, 10 mM NaCl, 3 mM MgCl2, 0.1% NP40, 1% BSA, 1 mM DTT, in the presence of RNase inhibitor (Roche 3335399001). After permeabilization cells were washed with 10 mM Tris-HCl pH 7.4, 10 mM NaCl, 3 mM MgCl2, 1mM DTT, 1% BSA with RNase inhibitor before being resuspended in 1× Nuclei buffer (10x Genomics) and counted. The sample was submitted to the CRUK Cambridge Institute Genomics Core for library preparation and 10X Genomics multiome using Chromium Next GEM single-cell multiome ATAC and gene expression reagent kit (10X Genomics, PN-1000283) according to the manufacturer’s instructions. Illumina sequencing was performed by the CRUK Genomics Core Sequencing facility on a NovaSeq X.

#### Reference genome

We downloaded gene annotation (.gtf) and DNA sequence (.fa) files from https://www.ensembl.org/. To prepare the reference genome for alignment, we used the mkgtf command from Cell Ranger [81] to filter out non-coding-protein genes from the raw .gtf file. The filtered .gtf and the genome .fa files were then used as input for the mkref command to generate the reference package compatible for alignment by Cell Ranger. In this manuscript, we have built reference packages for human (*Homo Sapiens*, GRCh38) and rat (*Rattus Norvegicus*, mRatBN7).

The rat genome is known to contain a large Numt region on chromosome 1 which can affect alignment to the mitochondrial genome [82]. We mask the Numt regions to prevent any mitochondrial reads from being aligned to the chromosome. We use a .bed file containing the starting and ending position of the selected chromosomes that need to be masked. Practically, we used maskfasta command from bedtools (v.2.18) to modify the genome .fa in the masked regions.

#### Single-cell data processing

We used 10x Genomics Cell Ranger software (v.7.2) to process raw scRNA-seq data. Specifically, we applied the count command to align sequencing reads to the reference genome and generate the aligned reads in .bam and the gene expression matrix based on unique molecular identifier (UMI) count in .tsv for each sample after merging multiple runs for the same library. For multi-modal datasets that include V(D)J and cell surface proteins, we used the multi command to also run the V(D)J pipeline and generate clonotype information in .tsv files.

Data from Terekhova et al. [34] is multiplexed-each sequencing library was prepared over pools of 6 samples with 2 technical replicates each. Briefly, the authors perform de-multiplexing using a combination of genotype-based demultiplexing using the souporcell pipeline and hashtag-based demultiplexing using HTODemux in Seurat. We verified the demultiplexing approach for one pool of samples using publicly available code. We used the author-provided barcodes to demultiplex all the pools.

We then performed a quality control on the aggregated gene expression matrix using Scanpy (v.1.7.2). To filter out potential doublets or multiplets, we discarded cells with more than 3000 detected genes or more than 10,000 total UMI counts. Additionally, to remove lysed or stressed cells, we filtered out cells with over 10% of reads mapping to the mitochondrial genome.

#### Cell annotations

When available in the original paper, we reused the provided cell annotations for our analysis. For datasets lacking the cell annotations, we inferred cell states or cell types using the gene expression matrix in combination with known marker genes. For each dataset, we took the aggregated gene expression matrix to perform the Leiden clustering using Scanpy (v.1.7.2), sweeping the resolution parameters to identify robust clustering. We chose a preferred resolution value based on cluster stability, such as a consistent number of clusters across the neighbouring resolutions. For each cluster *i, i* ∈ [0, 1, · · · *, n_c_*], we performed differential gene expression (DGE) analysis against all other cells. This yielded a list of genes differentially expressed in cluster *i*, potentially representing cluster-specific marker genes.

We downloaded a list of marker genes for each cell type across various tissue types in human and rats from Cell Marker 2.0 (http://bio-bigdata.hrbmu.edu.cn/CellMarker/). Given the high variability in the number of marker genes per cell type, we selected the top 10 genes with the highest number of supporting literature references for each cell type. For each cluster *i*, we computed the percentage overlap between the differentially expressed genes (DEGs) and the marker genes for all plausible cell types. A cluster was typically annotated as a specific cell type if more than 50% of its DEGs overlapped with that cell type’s marker genes.

A known limitation of this approach is that a single cluster may be assigned to multiple cell types, particularly when those cell types share marker genes (e.g., T cells and NK cells, or monocytes and macrophages). To resolve these ambiguous assignments, we examined the expression levels of marker genes unique to one cell type. High expression of the unique marker gene favoured annotation to that specific cell type over others. We verified our annotations using reference-based cell typing via SingleR (v.2.10.0, [83])

#### mtDNA variant calling

We defined mtDNA variants or mutations as mismatches between RNA reads and the reference genome. We used the in-house pipeline first developed by Green et al. [26] to detect variants on cells passing the quality control on the UMI and feature counts [27]. The pipeline processes the .bam files for each individual and subsets the file to only include mitochondrial reads using samtools (v.1.19). Given the cell barcodes of good-quality cells, we used subset-bam (v.1.1.0) to split the mitochondrial .bam to be cellular specific. Then, we attempted to find mismatches between the reads and the reference genome for each cellular .bam file using pysam (v.0.22.0).

We use a consensus approach for variant calling with UMIs, every base is assigned the majority of the UMIs, and it is reference if there are an equal number. Taking read consensus while variant calling exponentially decreases the chance that a random sequencing error on a read survives the final consensus call. We further filter out low-confidence genome sites, default any ties to the reference to avoid over-calling of variants, and mask read ends to minimize misalignment artifacts. To confidently call for mutations, we only considered callable sites in the mitochondrial genome which have at least 10 aligned reads, each having a base quality score above 25. We performed additional depth-based quality control to remove poor-quality cells with fewer than 100 callable sites. We also applied a 5-bp flanking filter to avoid false-positive variants near the end of the reads. Once mutations were called, we calculated heteroplasmy for each mutation as the proportion of reads carrying the mutation relative to the total number of reads aligned to that position. To allow multiple mutations in the same position, we defined heteroplasmy (*h_i_*) of a base *i* ∈ [*A, G, C, T*] with *N_i_* number of reads of base *i* at that position as:

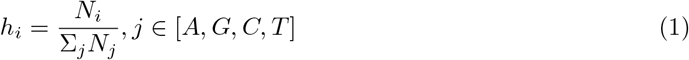

By the given definition, heteroplasmy *h_i_* acts as a proxy to the true proportion of mutated mtDNA in a single cell. To further avoid contamination from the PCR amplification stage, we excluded singleton mutations (i.e., mutations observed in only one cell) with heteroplasmy *h_i_ <* 0.3. We also excluded non-singleton mutations whose maximum heteroplasmy across all cells carrying the mutations was below 0.3.

For each mutation, we also defined its prevalence *p_i_* as the proportion of cells carrying the mutation among all cells possible to detect a mutation (i.e. at least 10 reads at that position and with at least 100 callable sites for that cell). We considered mutations with *p_i_* ≤ 0.01 as rare somatic mutations, denoted as low prevalence (LP) mutations. To ensure the reliability of the prevalence *p_i_*, we only included mutations that were callable in more than 100 cells in that position.

#### Mutant classification

To classify mtDNA mutations by their functional changes in the human genome, we used HmtVar [84] to further distinguish amino acid-altering mutations (non-synonymous, NS) from those that do not change the protein sequence (synonymous, S). We constructed the dictionary object in Python for each organism that consists of a pair of mutation IDs and the class of mutations (S or NS). For mutations occurring in non-coding protein genes, we classify the mutation as NaN. For NS mutations, we used MutPred [85] to further classify based on the pathogenicity score. We used a standard cut-off point to define high and low-pathogenic NS mutations.

We also classified mtDNA mutations based on the nucleotide changes as transitions and transversions. Transitions are the mutations in which a purine (A and G) and a pyrimidine (C and T) base change to the other purine and pyrimidine, respectively. Whereas, transversion is a change from purine to pyrimidine class, and vice versa.

#### CLP load

On average, 10x scRNA-seq data enables the observation of mutations at only ∼10-20% of the mitochondrial genome (see Supplementary S1), constrained to callable sites–those backed up by at least ten unique UMI reads and observed in over 100 cells. To estimate the total mutational burden across the mitochondrial genome, we calculated the frequency of CLP mutations among callable sites and scaled it up to obtain a genome-wide measure, which we term CLP load.

For a given donor *d*, we denote the number of unique CLP mutations observed across all aggregated cells of a specific type as *x_d_*, and the total number of callable bases across cells as *B_d_*. We model the total number of observed CLPs across donors as a Binomial process with mutation frequency *p*:

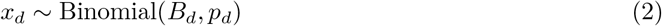

Assuming a Beta prior for *p*, we employed MCMC sampling to estimate its posterior distribution. To avoid overcounting, we sub-sampled cells sharing a common CLP mutation (see Supplementary S5.3), ensuring the independence of the number of unique mutations of each cell.

The estimated mutation frequency *p_d_* can then be extrapolated to the entire mitochondrial genome (∼16,000 bp) to obtain the expected number of CLP mutations per cell for donor *d* – CLP load. Conceptually, this is analogous to estimating the number of black balls (mutated sites) in a bucket of ∼ 16k of mostly white balls (non-mutated sites). The inference of *p_d_* in Eq.(2) and scale it up to ∼ 16k allows the estimation of true number of black balls given observing only ∼ 10% of the balls. By principle, CLP load is a reliable estimate if we assume that the frequency of mutations *p_d_* is uniform across the entire mitochondrial genome, ignoring a possible gene-specific variation in the mutation rates.

#### Selective pressure inferred from dN/dS

Non-synonymous (NS) mutations alter the amino acid sequence of proteins and may impact cellular fitness, making them subject to either positive or negative selection. To assess the selective pressure acting on NS mutations, we examined mtDNA protein-coding mutations and estimated the relative abundance of NS compared to synonymous (S) mutations. Assuming that S mutations are largely neutral and expand without selective constraints, and excess of NS mutations relative to this neutral baseline would indicate positive selection, whereas a deficit would suggest negative (purifying) selection.

In our analysis, we counted the number of unique NS and S mutations, denoted as *x_NS_* and *x_S_*, from the total number of possible sites *B_NS_* and *B_S_*, respectively. Treating these counts as outcomes of independent Binomial processes, we inferred the underlying mutation frequencies *p_NS_* and *p_S_* using pMCMC sampling. We defined the mutational bias 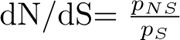 as a measure of selective pressure:

- dN/dS*>* 1 indicates positive selection,
- dN/dS*<* 1 indicates purifying selection, and
- dN/dS≈ 1 suggests neutrality.

To assess statistical significance, we computed 95% credible intervals (displayed in whiskers), and evaluated whether the neutral expectation (dN/dS= 1) falls within them. Intuitively, better mitochondrial genome coverage results in narrower credible intervals that improves confidence in the inference.

Although it is possible to compute dN/dS for each donor individually, in practice, many samples are underpowered due to lack of observed NS or S mutations. This is expected, as some mtDNA mutations occur in non-coding regions, which do not contribute to either NS or S. Moreover, the human mitochondrial genome is known to exhibit roughly a 3:1 ratio of NS to S mutations, further increasing the chance of samples no observed S mutations. To overcome this, we aggregated cells across donors having the same cell-type and tissue identity. We applied this method primarily to cryptic or low-prevalence (CLP) class, as well as to maximally prevalence mutations as comparison.

#### The coalescent model

A.G., F.I., P.F.C., and N.S.J. are funded by the Wellcome Collaborative Award (224486/Z/21/Z). N.S.J. is funded by Leverhulme (RP G 2018 408). P.F.C. is currently funded by a Wellcome Discovery Award (226653/Z/22/Z), the Leverhulme Trust (RP G 2018 408), the Medical Research Council Mitochondrial Biology Unit (M C U U 00028/7), and the Biological and Biotechnology Research Council (BB/Y 003209/1), the Rosetrees Trust (P GL23/100048), and the LifeArc Centre to Treat Mitochondrial Diseases (LAC-TreatMito) under grant no. 10748. LifeArc is a charity registered in England and Wales under no. 1015243 and in Scotland under no. SC037861. His research is supported by the NIHR Cambridge Biomedical Research Centre (BRC 1215 20014). The views expressed are those of the author(s) and not necessarily those of the NIHR or the Department of Health and Social Care. K.S. is a recipient of a scholarship from the Indonesia Endowment Fund for Education (LPDP), Ministry of Finance of

To characterise how the heteroplasmy distribution evolves with age, we employed a truncated coalescent model that represents the backward-in-time analogue of the Moran process. Specifically, for a sample of *n* mtDNA molecules drawn from a total of *N*, we traced the genealogy backward in time up to the mitochondrial age *W*, recording the branch lengths with exactly *b* descendants, denoted by *L_b,n_, b* ∈ {1, 2*, …, n*}. These branch lengths determine the shape of the heteroplasmy distribution. Then, we assumed the number of somatic mutations at heteroplasmy *b/n*, denoted by *m_b,n_*, follows a Poisson distribution with mean *θ* · *L_b,n_*, where *θ* represents the mutation rate. Intuitively, younger individuals are expected to have a smaller mitochondrial age *W* and are less likely to have reached the most recent common ancestor (TMRCA), resulting in fewer or no homoplasmic mutations. In contrast, older individuals with a larger *W* are more likely to accumulate such mutations (see Supplementary S5 for schematic figures and a more detailed explanation adapted from Green et al. [26]).

We fit this model to human and rat data using Bayesian inference, estimating the individual-specific mitochondrial age *W* and mutation rate *θ*. Notably, both parameters are affected by known confounders: *W* = *α* · *τ*, where *τ* is chronological age, and *θ* = *B* · Θ, where *B* is the average bases passing per cell. Hence, we instead inferred:

- the ageing rate *α*, representing the change in mitochondrial age per year, and
- the scaled mutation rate Θ.

#### Differential Gene Expression analysis

Through our Differential Gene Expression (DGE) analysis, we compare cells containing at least one non-synonymous CLP mutation (heteroplasmy above 30%) to cells with no observed CLP mutations in a non-parametric manner. This analysis is performed over T cells from 3 datasets that satisfy our sample size constraints [34, 35, 38]. We combine cells across all donors for statistical power. After preprocessing for single-cell sequencing quality, we normalize the data using counts per 10k and perform the DGE over normalized data.

Cells harboring a CLP mtDNA mutation are expected to have a higher coverage across the mitochondrial genome (higher number of sites covered). Since somatic CLP mtDNA mutation levels increase with age, cells harboring a CLP mtDNA mutation will have a higher donor-age distribution compared to cells from donors without a CLP mtDNA mutation. To robustly ascribe the results of the DGE to the presence or absence of CLP, we need to control for donor age, and the total sites covered for the two cell populations we want to compare using DGE.

Therefore, we match the distribution of cells without a CLP mtDNA mutation to the distribution of cells with at least one CLP mutation across total mitochondrial sites covered and age. We use a two dimensional histogram to sample from the joint distribution of total sites and age. We use importance sampling to sub-sample the cells without a CLP mtDNA mutation, in a similar fashion to Coarsened Exact Matching. The size of the sub-sampled set is determined based on the histogram overlaps and computing the standardized mean difference (SMD). We pick the largest sub-sample such that the histogram overlap of the marginals is above 0.8 and the per-feature SMD (Cohen’s d) is lower than 0.2.

We repeat the sub-sampling and testing procedure 100 times. We perform a Mann-Whitney U test comparing the set of cells with a CLP mutation with each subsample. For each sub-sampled DGE run, we obtain a list of significant genes after multiple hypothesis correction (*p_adj_ <* 0.05, Benjamini-Hochberg correction). To robustly identify the genes indicative of biological signals - we look at recurring genes that are significant in more than 30% of the sampling runs and use them for further analysis. Further details of the subsampling procedure can be found in Supplementary S6.

#### Gene Ontology analysis

The measured gene expression per-cell depends on a multitude of factors - such as sequencing methodology, cell state, and tissue type. However, it is possible for the same cell type to express different genes linked to similar biological pathways at different times. We perform a Gene Ontology Overrepresentation analysis per dataset, to statistically interpret the pathways each set of recurring, significant genes is enriched for using the Enrichr API via the *gseapy* python package, queried over the C5 ontology gene sets collection (c5.go.bp.v2025.1.Hs.symbols.gmt) from the Molecular Signatures Database [86–89]. We report significant GO terms with *p_adj_ <* 0.05.

#### Individual-level meta-analysis

We perform an individual-level random-effects meta-analysis procedure using PyMARE over three T cell datasets [34, 35, 38] that satisfy our sample size constraints. For each gene, for each individual— we perform 100 Mann-Whitney U tests comparing subsampled cells with Non-Synonymous CLP mutations and cells with No CLP mutations, obtaining a distribution of area under the curve (AUC) estimates. The sample mean and sample variance of this sampled AUC distribution serves as the effect size estimate and its sampling variance for the random-effects meta-analysis. We transform the AUC to its logit scale to satisfy normality assumptions for the meta-analysis. Individual-level logit(AUC) estimates are combined using a variance-based likelihood estimator with maximum likelihood optimization. Gene-level significance is measured via a Z-test and we use the Benjamini-Hochberg method to correct the p-values for multiple testing across genes. Full gene lists can be found in the Supplement. All the genes, along with p-values, are passed to scPPIN for visualizing the gene interaction network (https://floklimm.shinyapps.io/scPPIN-online/) [90].

## Data Availability

The publicly available scRNA-seq datasets analysed in this study were obtained from the Gene Expression Omnibus under accession numbers GSE157007, GSE136184, GSE120221, GSE135194, GSE133181, GSE136831, GSE260769, GSE130973, GSE143704, and GSE137869; Synapse under accession syn49637038; the Genome Sequence Archive for Human under accession HRA000395; and BioStudies under accession E-MTAB-13874. Further dataset details are provided in Supplementary Table 1. The newly generated scATAC-seq data are currently being deposited in a public repository, and their accession details will be provided once deposition is complete.

## Supporting information

Supplementary Tables for Fig 4.

## Acknowledgement

Cell sorting/flow cytometry analysis for this project was done by R. Schulte and G. Grondys-Kotarba from the Cambridge Institute for Medical Research Flow Cytometry Facility for cell sorting. The CRUK Cambridge Institute Genomics Core Facility for library preparation and 10X Genomics multiome and sequencing services. We thank Dr. Maria Grazia Spillantini and Dr. Emre Fertan for discussions on the manuscript.

## Funding

A.G., F.I., P.F.C., and N.S.J. are funded by the Wellcome Collaborative Award (224486*/Z/*21*/Z*). N.S.J. is funded by Leverhulme (*RPG* − 2018 − 408). P.F.C. is currently funded by a Wellcome Discovery Award (226653*/Z/*22*/Z*), the Leverhulme Trust (*RPG* − 2018 − 408), the Medical Research Council Mitochondrial Biology Unit (*MC UU* 00028*/*7), and the Biological and Biotechnology Research Council (*BB/Y* 003209*/*1), the Rosetrees Trust (*PGL*23*/*100048), and the LifeArc Centre to Treat Mitochondrial Diseases (LAC-TreatMito) under grant no. 10748. LifeArc is a charity registered in England and Wales under no. 1015243 and in Scotland under no. SC037861. His research is supported by the NIHR Cambridge Biomedical Research Centre (*BRC* − 1215 − 20014). The views expressed are those of the author(s) and not necessarily those of the NIHR or the Department of Health and Social Care. K.S. is a recipient of a scholarship from the Indonesia Endowment Fund for Education (LPDP), Ministry of Finance of the Republic of Indonesia. H.J. is supported by EPSRC (EP/S023151/1).

## Supplementary Discussion

## S1 scRNAseq preprocessing and variant calling

### Robustness to additive noise

Any mtDNA variant obtained from the pipeline can be noisy: the mutation can either be a true mutation with a heteroplasmy estimation error, or it can be a mistakenly called non-existent mutation i.e. a false positive. We apply strict filtering thresholds to minimize false positives (see Methods), but random errors can still lead to additive noise across variants. Our downstream analyses depend on comparing heteroplasmy distributions of variants across age groups, rather than absolute estimates from individual variant calls. We would expect any age-independent additive noise to blur age-associated distributional shifts but in our results we still observe a strong age-associated signal implying that the effect of any additive noise is limited. Aggregating across cells with CLP variants further minimizes the effect of additive noise. We also corroborate our analysis by reproducing the main results using variants obtained from newly-generated mt-scATAC-seq data.

### Mitochondrial coverage by sequencing modality

**Fig. S1.1.**
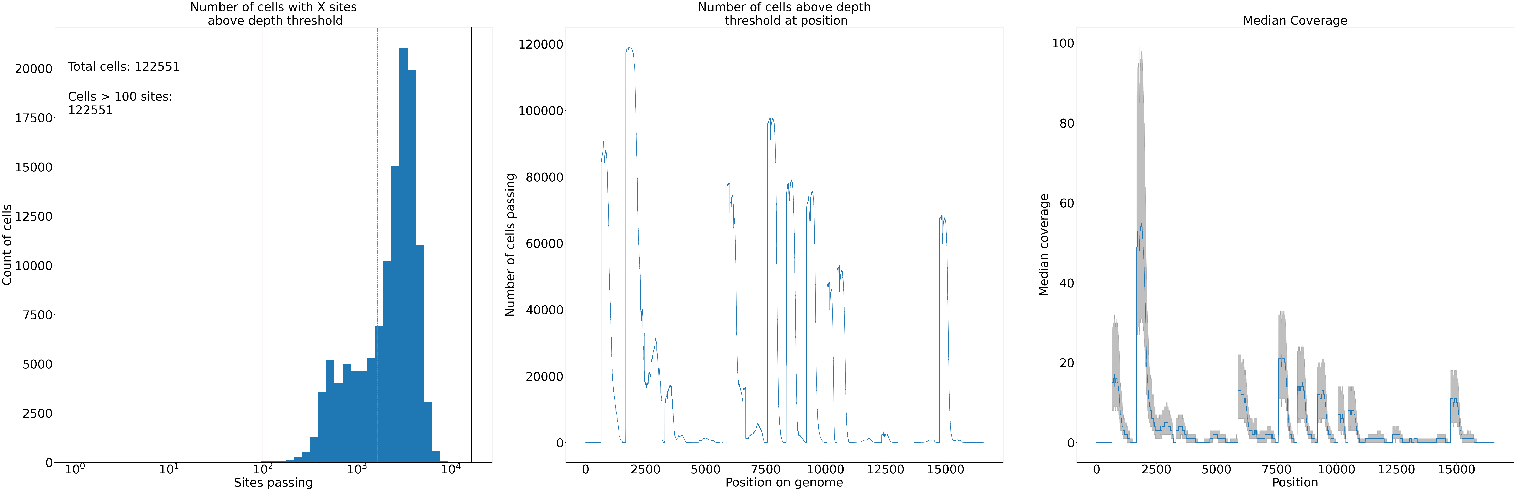
Illustrating depth and breadth of mtDNA coverage for 10x 5’ scRNA-seq data from Luo et al. [35]

**Fig. S1.2.**
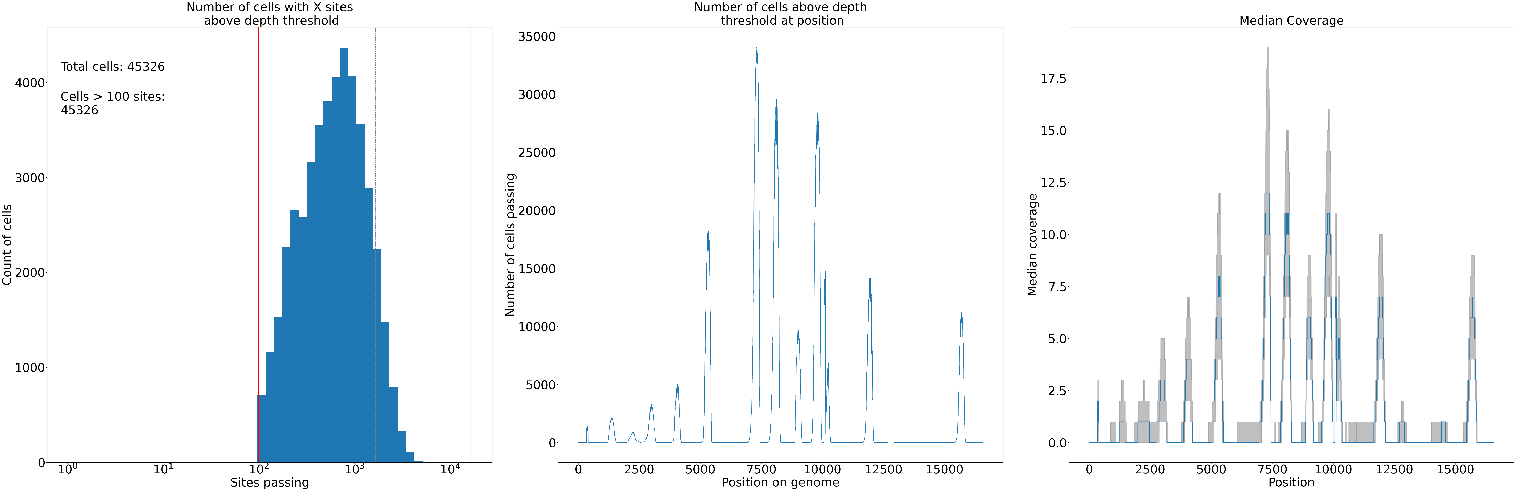
Illustrating depth and breadth of mtDNA coverage for 10x 3’ scRNA-seq data from Oetjen et al. [39]

**Fig. S1.3.**
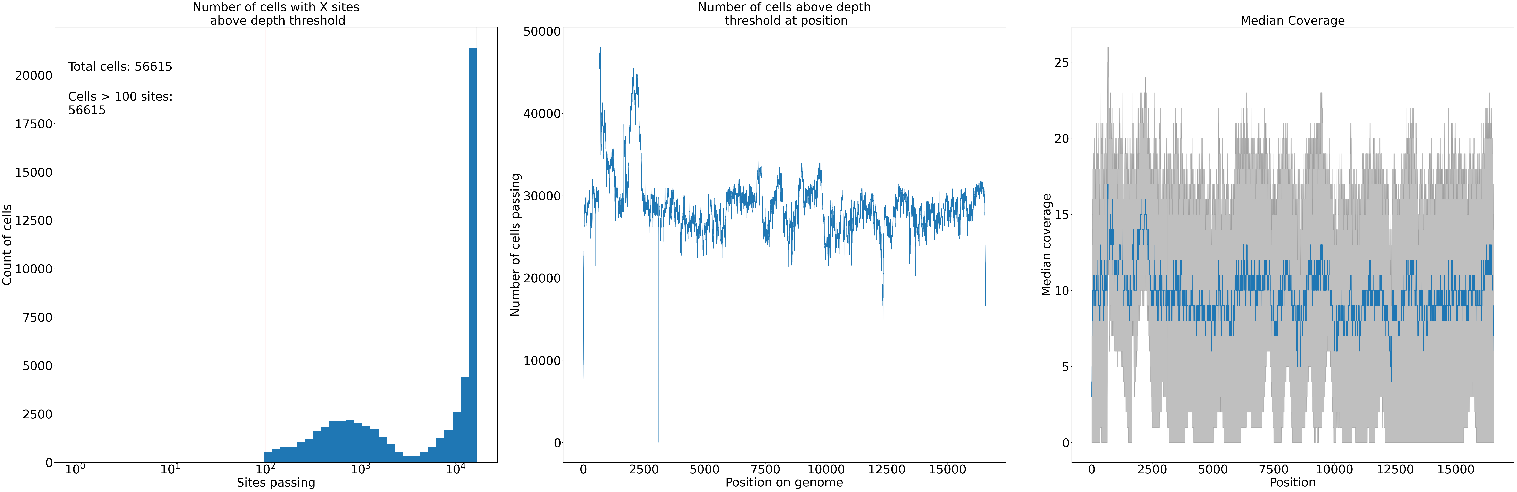
Illustrating depth and breadth of mtDNA coverage for newly generated mt-scATAC-seq data

## S2 Age-dependent dynamics of CLP mtDNA mutations

### S2.1 Distribution of the number of CLPs per cell by mid-late life

**Fig. S2.1.**
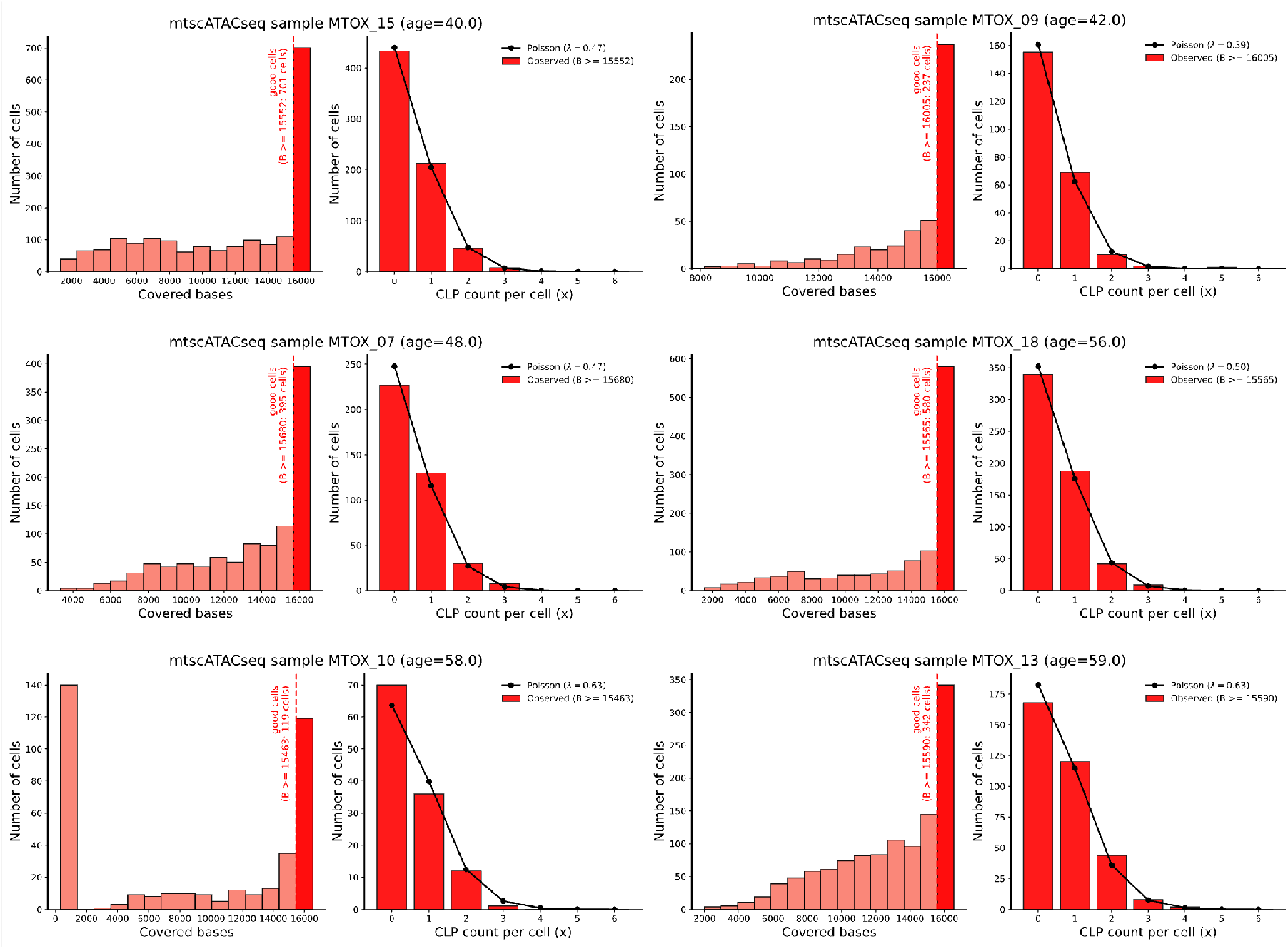
We estimate that ∼40% of cells have at least one CLP by mid-late life. From the newly generated mtscATAC-seq data, we first inspect the number of covered bases across T cells (left panel) from six donors aged around 50 years and then select cells with at least 90% genome coverage to examine the distribution of CLPs per cell (right panel). The observed CLP counts follow a Poisson distribution with rate ∼0.5 CLPs per cell. This implies that, by mid-late life, although the average CLP burden is ∼0.5 per cell, only about 40% of cells are expected to carry at least one CLP.

### S2.2 Abundance of CLPs and distribution of CLP clone sizes

**Fig. S2.2.**
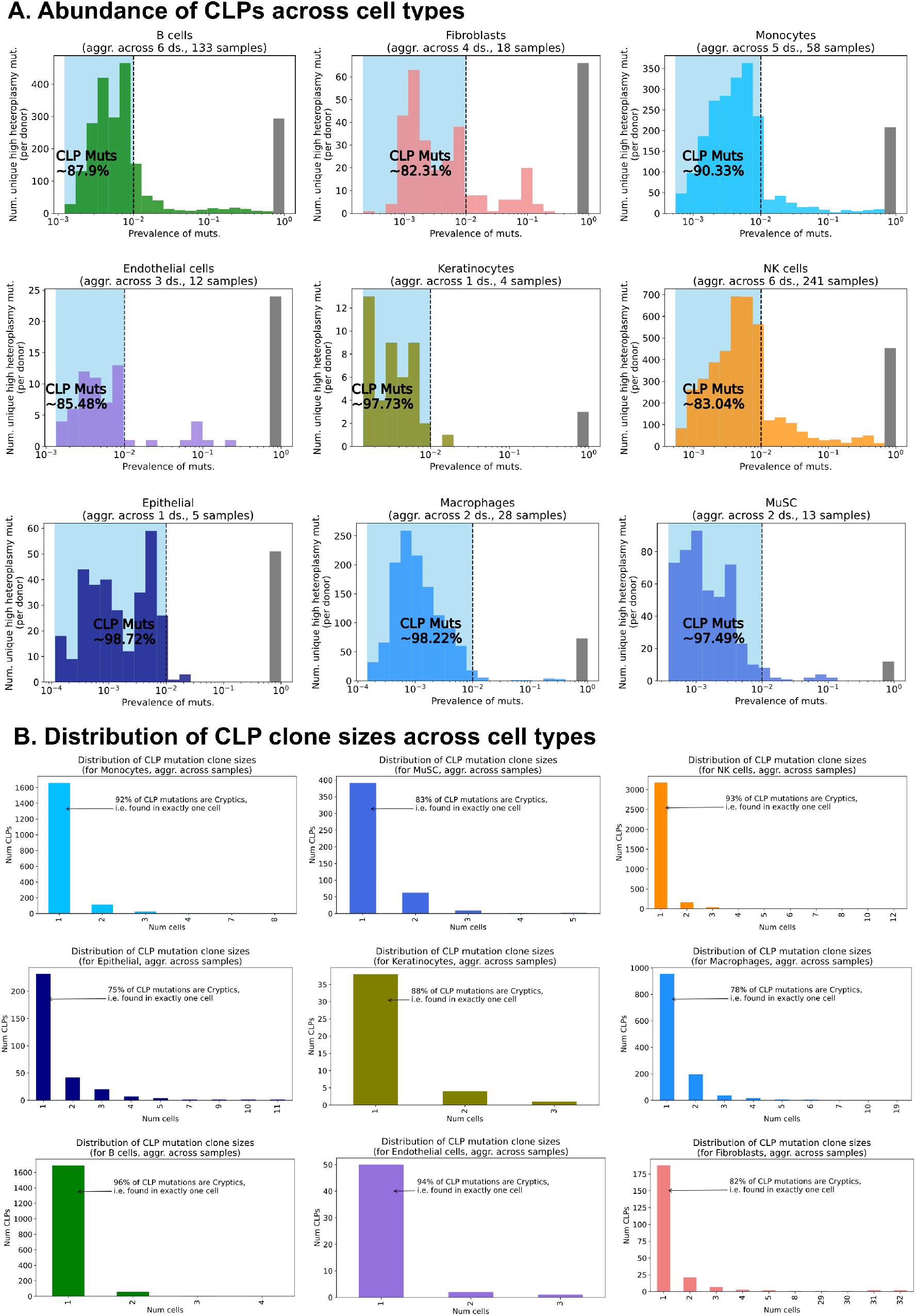
CLPs are abundant across cell types and most CLPs are consistently cryptic, i.e. unique to a single cell. Subpanels A and B are extensions to Fig 1C and Fig 1D respectively.

### S2.3 Heteroplasmy distributions with age in human proliferative cells

**Fig. S2.3.**
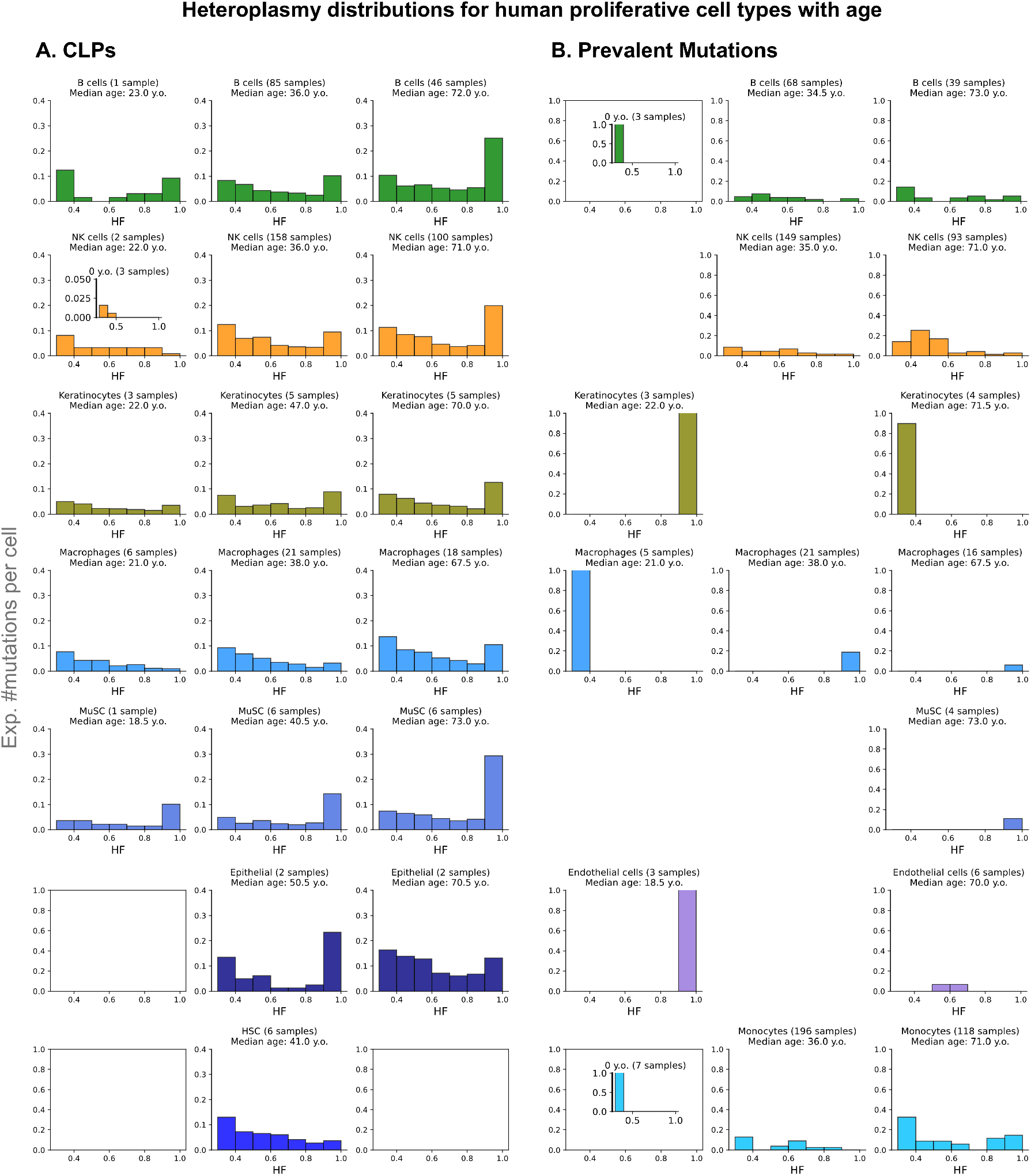
Heteroplasmy distributions of CLPs consistently shift to higher heteroplasmies and homoplasmies with age, unlike more prevalent mutations, across cell types and tissues. The heteroplasmy distributions shown here are across 7 cell types, have been generated after clonal deduplication, and are normalised by total genome sites passing, scaled to the entire genome. This is an extension to Figure 1D.

### S2.4 Covariation of CLP loads across human proliferative cell types

**Fig. S2.4.**
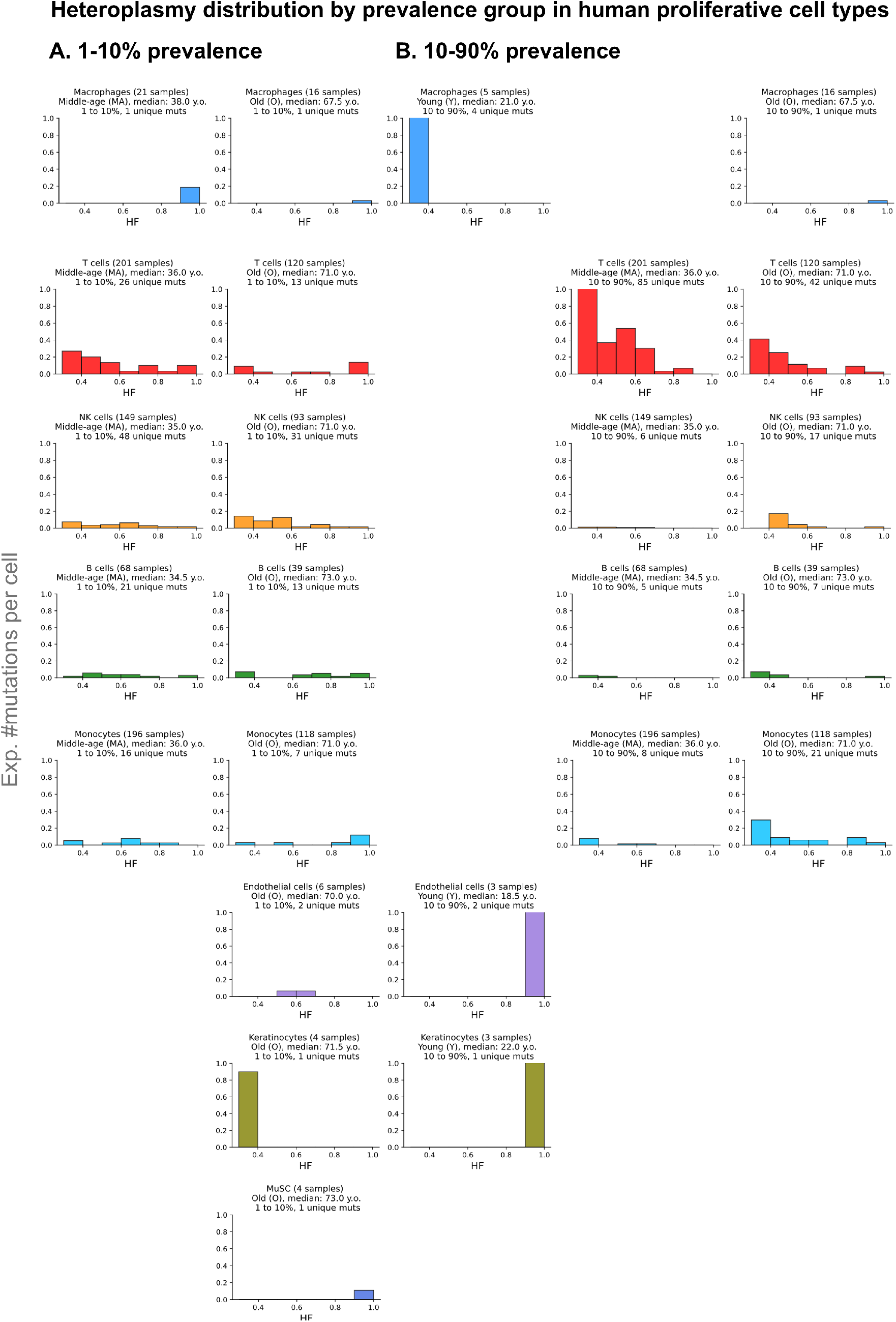
Limited evidence of age-associated accumulation in prevalent mutations; broken down by prevalence group, showing aggregate distributions for mutations with 1 − 10% prevalence (A) and 10 − 90% prevalence (B) respectively.

### S2.5 Comparing CLP load and chronological age

**Fig. S2.5.**
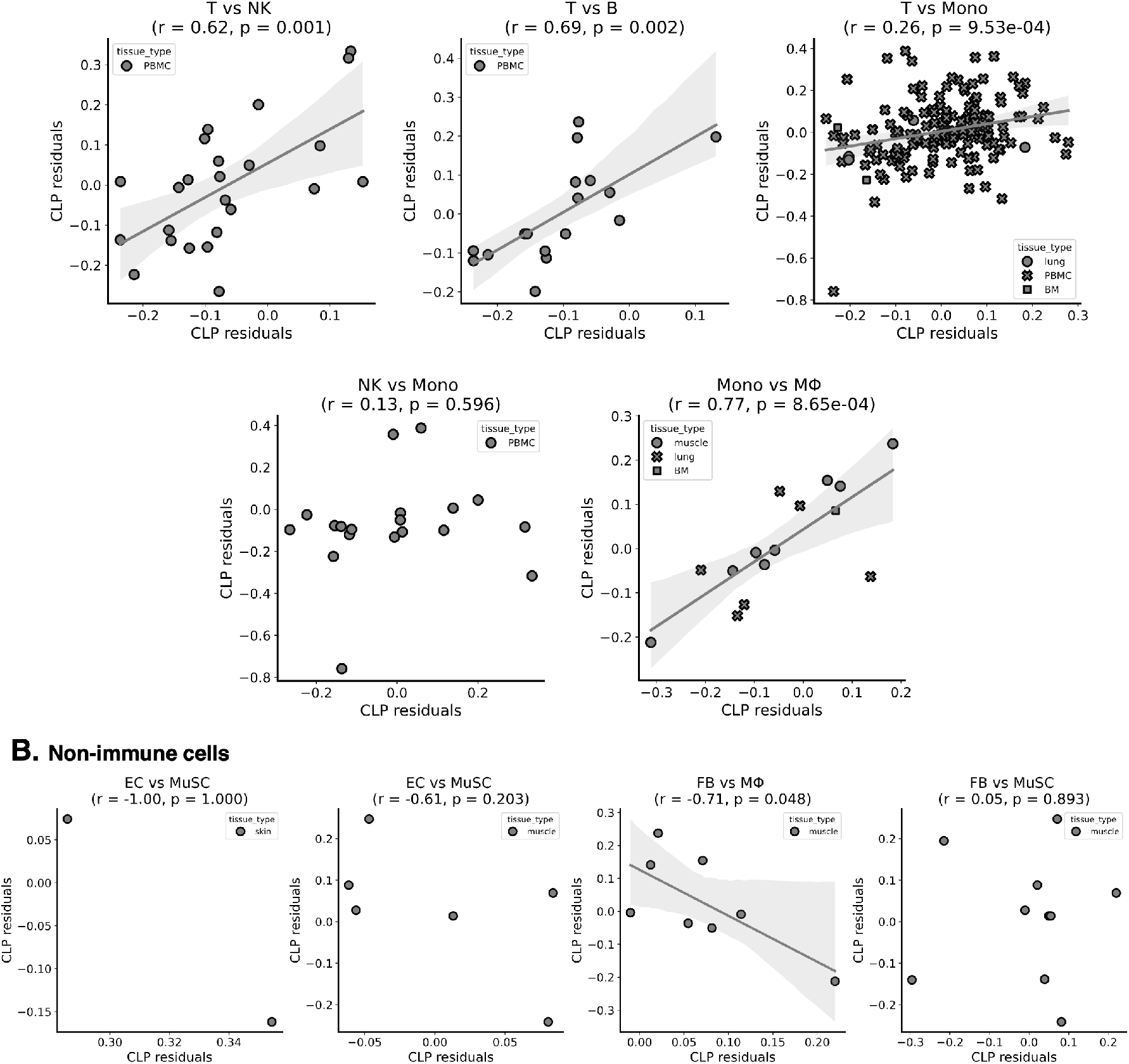
CLP residuals covary positively across immune cell types obtained from the same sample. CLP residuals are defined as the difference in the observed CLP load compared with the expected CLP load from the linear fits shown in Figure 2A. We show significant positive correlation of CLP residuals (Pearson-r and p-values) between different immune cell types belonging to the same sample, which might point to shared donor-level factors that might affect CLP accumulation. For non-immune cell types, we demonstrate a lack of relative statistical power to make similar claims. This is an extension to Figure 2D.

### S2.6 CLP mutations in rat proliferative cells

**Fig. S2.6.**
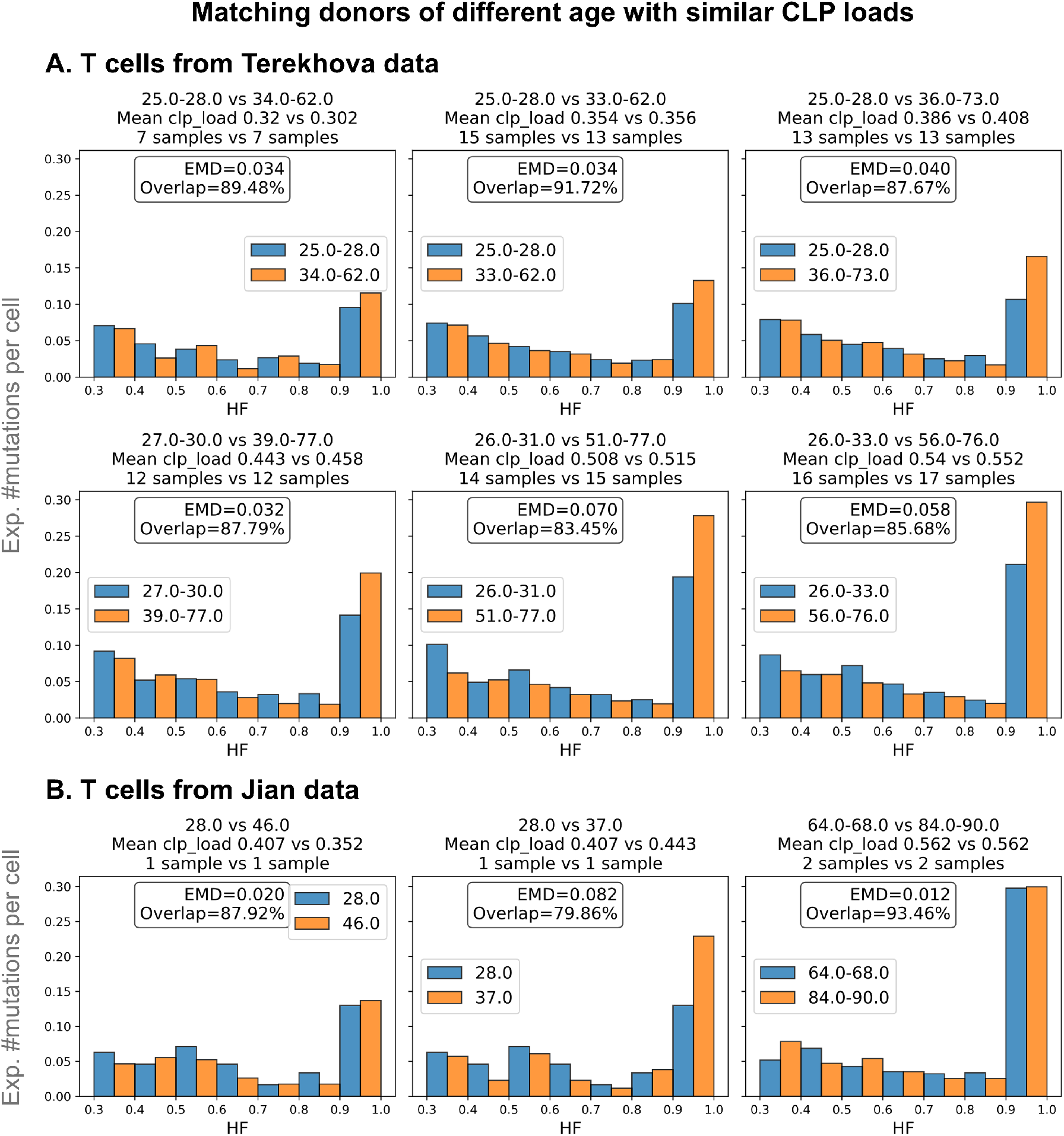
The heteroplasmy distributions of donors in different age groups with matched CLP loads are broadly similar, accounting for age-associated fixation. On matching by CLP load, the distributions for elder donors are similar to the distributions for the younger donors, with EMD values ranging between 0.012 − 0.082 across two different datasets. As expected, the distributions of older individuals have slightly more fixed homoplasmies, but both the elder and younger groups have qualitatively similar distributions, consistent with the dynamics expected from previously presented evidence of age-associated accumulation and of neutral selection with age (see Fig 2, Fig 3); *Method for matching CLP load:* For a given CLP load *C*, we find the subset of donors in a given dataset in the *C* ± 0.05 range, and compare the aggregate heteroplasmy distributions of the bottom and top 0.25 quartiles of this subset by chronological age. We sweep over *C* ∈ {0.30, 0.35*, …,* 0.55} and run the analysis for groups with at least 1 year of difference between the minimum age of the bottom quartile and the maximum age of the top quartile. We use the Wasserstein distance (or EMD) and histogram overlaps to measure similarity between the bottom and top quartile distributions by age.

**Fig. S2.7.**
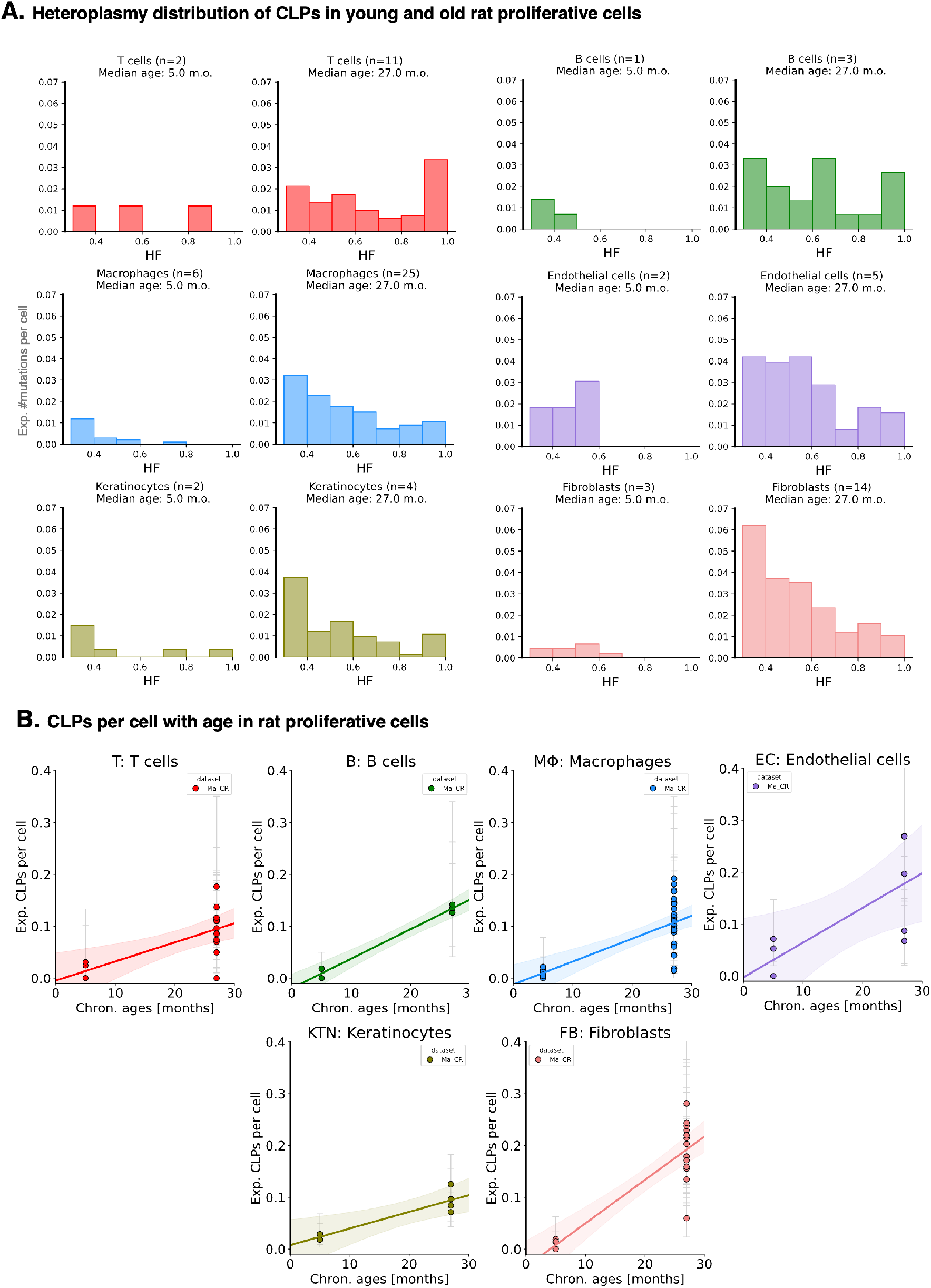
A. Cross-species validation of Figure 1D: Shifts in the heteroplasmy distribution of CLPs to higher heteroplasmy with age are also observed in rats from [91]; B. Cross-species validation of Figure 2A: We observe similar accumulation patterns to higher expected CLP load per cell with age in rats, although rat cells show higher per-annum accumulation rates: ∼10% cells have at least one CLP by age 24 months.

## S3 Polyclonality assumption

**Fig. S3.1.**
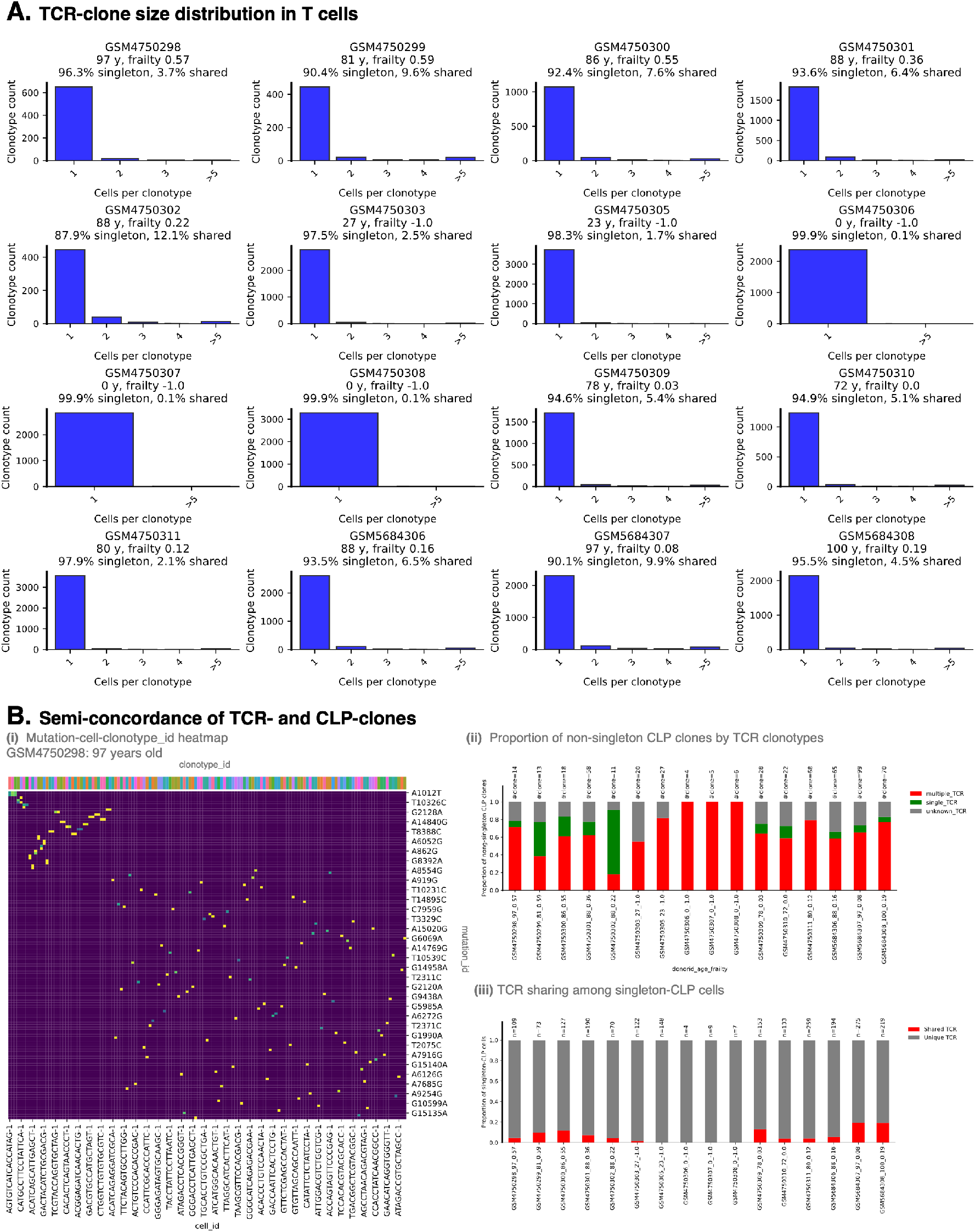
Most T cells are associated with unique TCR clones and CLP clones: **A.** We checked on 16 TCRseq data from Luo, et al. to identify clonotype identities for each T cell. Most TCRs represent small clones which are only observed once in the data, but in the rare event that clones have multiple cells in the dataset; **B(iii).** It is rare to see these cells having two or more different CLPs (multiple different mtDNA mutations per single TCR). In the rare cases where different CLPs are found within a single TCR the mtDNA mutations might actually be in all cells in each TCR but are not observed because of uneven coverage, or these could genuinely be mutations emerging within a single T cell receptor clone; **B(i-ii).** As we know in Fig.1 it is also rare for a particular CLP to be found in more than one cell. In the case that a CLP is in multiple cells, we see that the mutations can be shared across multiple TCRs, pointing to an origin of some mutations earlier in the cell hierarchy, e.g., among HSCs.

## S4 Selection across cell types and prevalence groups

**Fig. S4.1.**
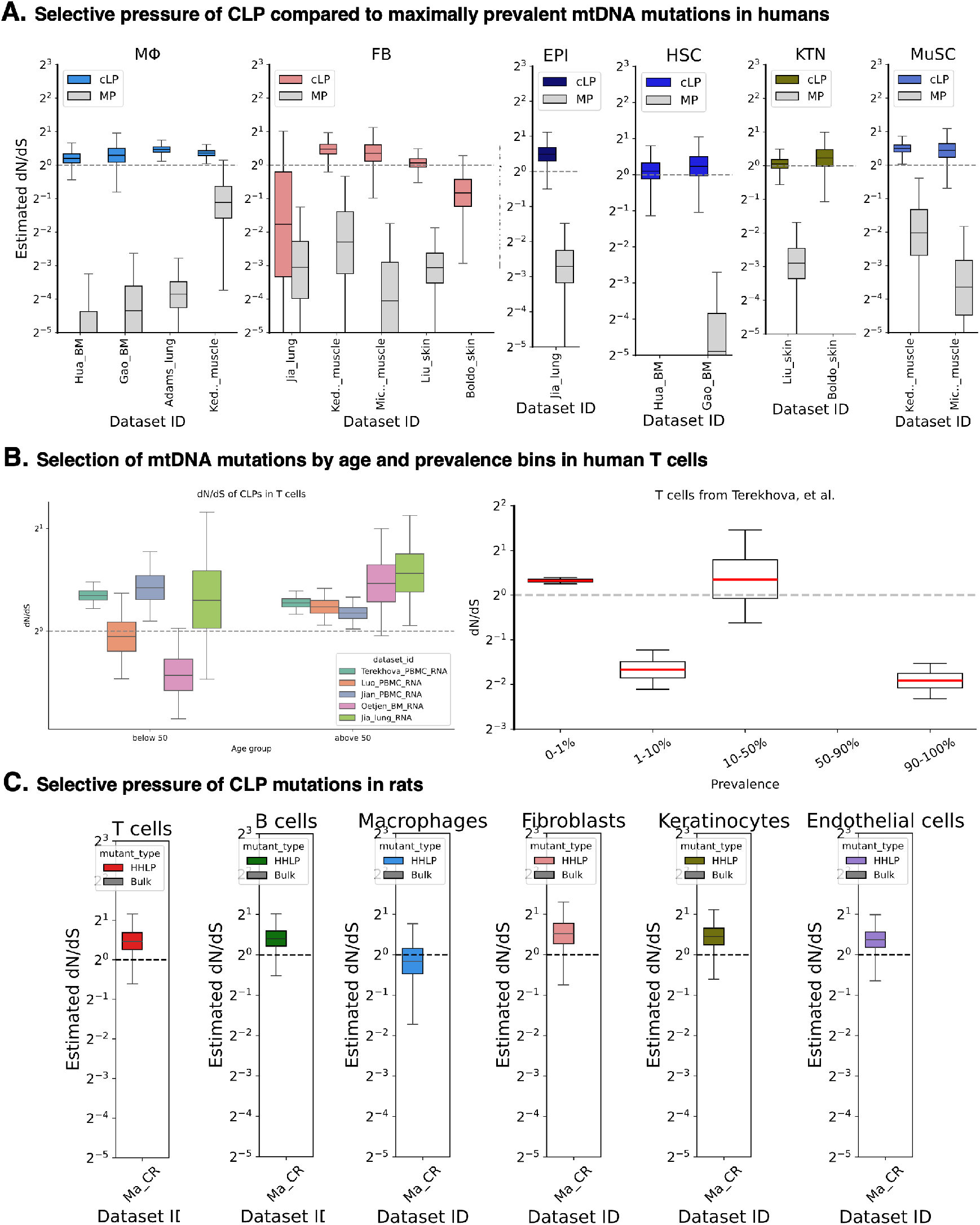
A. Estimated dN/dS ratios showing evidence of neutral or weak positive selection in Non-Synonymous CLPs across six cell types. Unlike CLPs, prevalent mutations, i.e. those with prevalence ≥ 1%, continue to be heavily selected negatively. This is an extension to Figure 3A; **B. Prevalent mutations are negatively selected when examined granularly, across 4 prevalence sub-groups**. We examine dN/dS ratios across prevalence bins for data from Terekhova et al. [34] et al. We see evidence of negative selection across prevalence groups 1 − 10% and 90 − 100%. For the 10 − 50% prevalence group, the relatively large confidence interval is due to a low number of mtDNA mutations detected in that bin. We see evidence of weak positive selection for CLPs (ie. the 0 − 1% prevalence group) consistent with Figure 3A. **C. Cross-species validation of selective pressure on rats:** We show CLP mutations continue to undergo neutral or weak positive selection across six cell types in rats, consistent with the observed patterns in humans in Figure 3A.

## S5 Parameter inference

### S5.1 Clonal deduplication by representative-cell subsampling

**Fig. S5.1.**
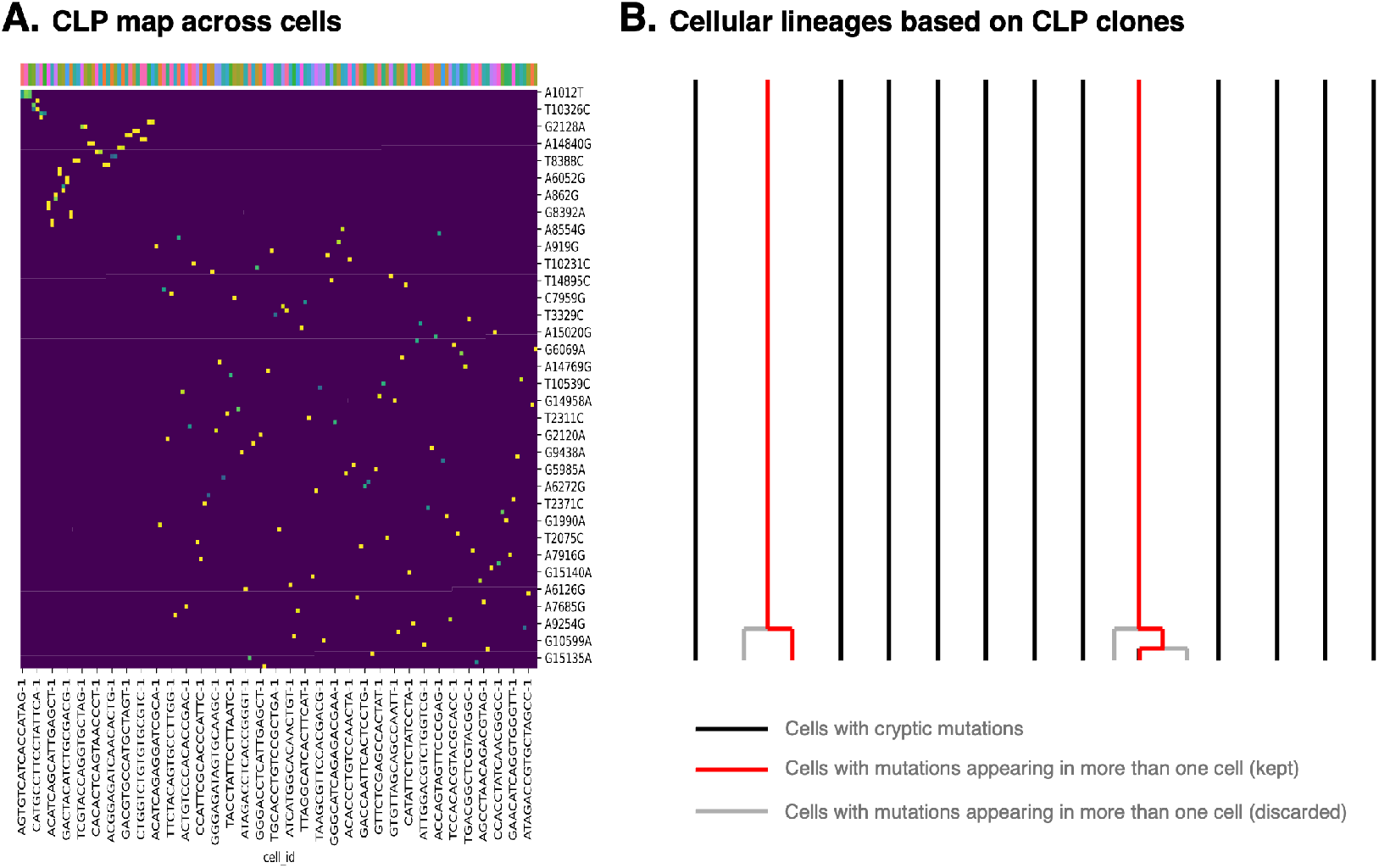
Shared-CLP subsampling to avoid double counting in the inference: **A.** Heatmap of CLP mutations detected across single cells in one example donor. Columns denote cell barcodes and rows denote mutation IDs with prevalence ≤1%. Cells with no detected mutations are omitted from the visualisation. Heatmap entries indicate the heteroplasmy of each mutation in each cell, with zero indicating that the mutation was not observed in that cell. Cells and mutations were hierarchically clustered according to their CLP-sharing patterns. Most detected mutations are unique to a single cell, whereas a small fraction of CLPs are observed in multiple cells, forming CLP clones; **B.** Most CLPs observed are unique to a single cell, hence most cell lineages are treated as independent (black vertical lines). In rare cases, one CLP is shared by more than one cell, forming a clone (red lineages). To avoid double counting in the inference, we randomly choose one representative cell from that CLP-clone and discard the rest (shown in grey). Hence, after subsampling, all retained CLP mutations are unique to a single cell. To match this subsampling, we also remove cells with no observed CLPs at the same rate as we remove cells with CLPs.

### S5.2 Truncated coalescent model and Bayesian inference

From the previous section, we find evidence that most cells arise from independent lineages, with only rare cases in which cells share a lineage. In other words, most cells in the samples appear to accumulate CLP mutations independently, similar to post-mitotic cells. To model the accumulation of somatic mtDNA mutations within a single cell, we use the truncated coalescent model proposed by Green et al. [26]. We argue that this model can also be reasonably applied to proliferative cells, given the polyclonal structure observed in our data.

Within a single cell, there are roughly 100-1000 mtDNA molecules that continually replicate and turn over. When an mtDNA molecule replicates, it may acquire a mutation, which can later segregate to higher heteroplasmy or be lost from the population. This forward-in-time process is well described by the Moran model [92]. The coalescent model is its backward-in-time equivalent.

The coalescent model is governed by two main parameters: *θ*, the scaled mutation rate, and *W*, the mitochondrial age. Given *θ* and *W*, we can predict the expected number of mutations *M_b/n_* that appear at heteroplasmy *b/n* in a single cell, and this framework can be extended to multiple cells. For younger individuals with lower mitochondrial age (Fig.S5.2A(i)), the lineage tree is dominated by external branches, so mutations tend to appear at lower heteroplasmy. In contrast, older individuals with higher mitochondrial age (Fig.S5.2A(ii)) have deeper coalescent trees, allowing mutations to reach higher heteroplasmy. Having access to mtDNA mutation data from scRNA-seq data allows us to perform parameter inference. Developed by Green et al. we attempt to infer donor-specific parameters, Θ*_d_* and *α_d_*, which are equivalents of *θ_d_* and *W_d_* respectively through *θ_d_* = Θ*_d_B_d_* and *W_d_* = *α_d_τ_d_*, with *B_d_* as the total sites observed and *τ_d_* as the chronological age of donor *d*. The full yet concise explanation of the model and inference technique is given as follows:

1. For the case of *n* samples (out of *N*), we evaluate *T^W^_k,n_* as the waiting time it takes for *k* lineages to coalesce to *k* − 1, given the mitochondrial age of *W*
2. Use *T^W^_k,n_* to evaluate the total length of branches that carry exactly *b* descendants, *b* = 1, 2, · · · *, n*, termed as *L^W^_b,n_* Any mutations occurring in these branches with total length of *L^W^_b,n_* will appear at *b/n* heteroplasmy, *b,n* The number of mutations that appear at *b/n* heteroplasmy
3. The number of mutations that appear at *b/n* heteroplasmy in a single cell, termed *M_b,n_*, follows Poisson distribution with the rate of *L^W^_b,n_*. Hence, we have:

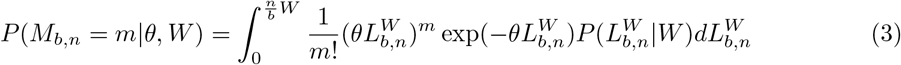

Which is intractable to integrate over all plausible branch lengths of the tree.
4. We can then extend Eq.3 for the case of a set of *C* cells carrying *M^C^_b,n_* each, giving the likelihood of data *D* to be:

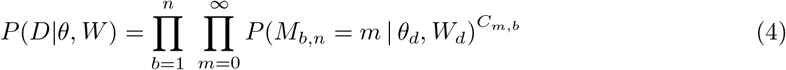

with *C_m,b_* is the number of cells carrying *m* mutations at heteroplasmy *b/n*,
5. In the current report, we used hierarchical Bayesian inference (Fig.S5.2B: each individual *d* has two target parameters Θ*_d_*and *α_d_*, which are sampled from population-level distribution *α_d_* ∼ Log-Normal(*µ_α_, σ_α_*) and Θ*_d_* ∼ Log-Normal(*µ*_Θ_*, σ*_Θ_). For a set of *K* donors with *C_d_* cells each, we have our final posterior as:

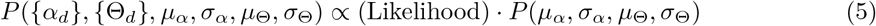

with the likelihood of

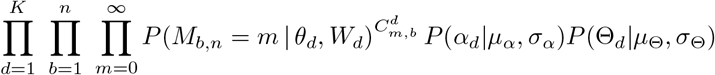

In this report, we use Monte Carlo to sample *L^W^_b,n_* and compute *E[L^W^_b,n_]* as well as perform a grid search to evaluate the posterior using Eq.5 over predefined parameter intervals.

**Fig. S5.2.**
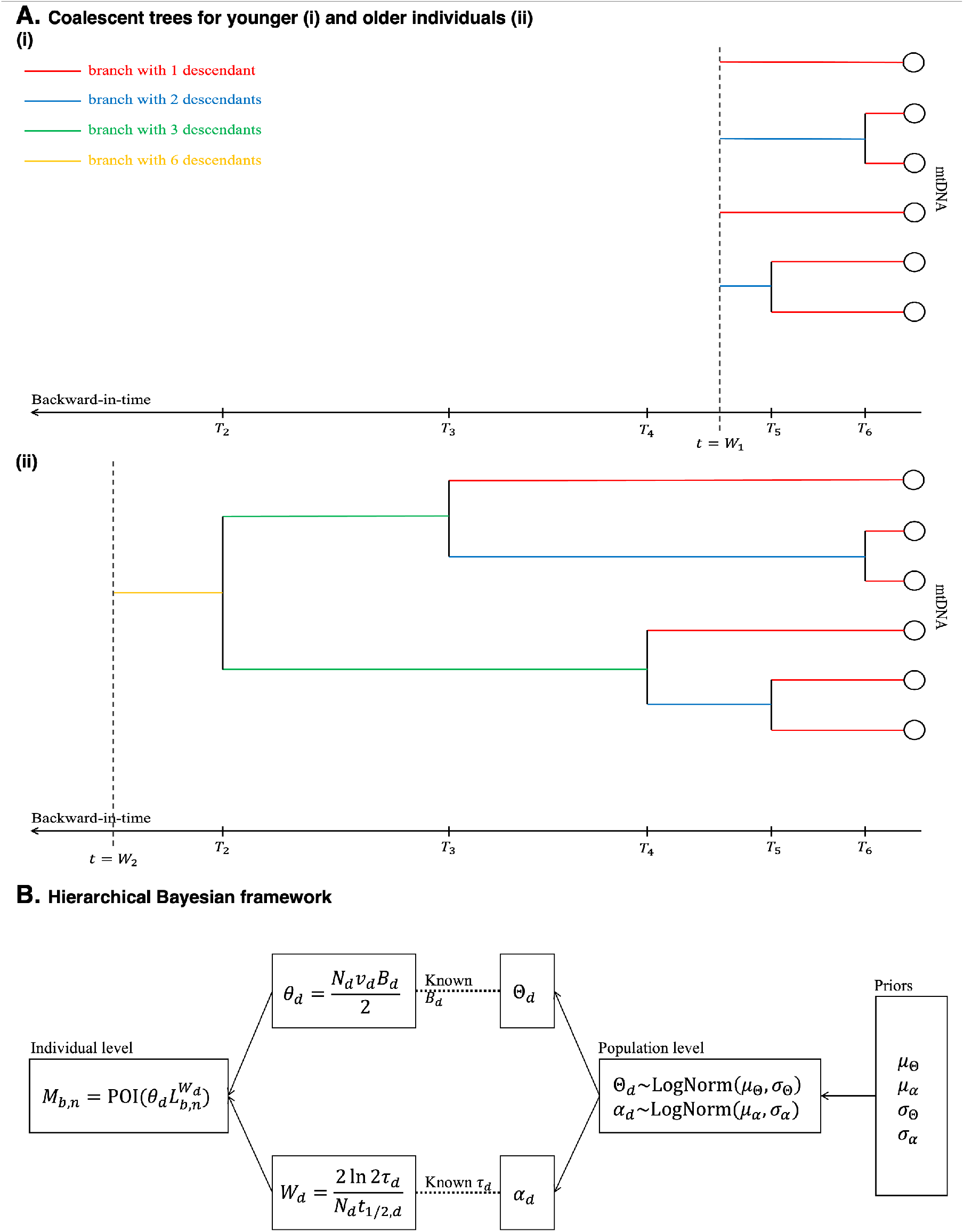
Truncated coalescent model for simulating CLP heteroplasmy distributions and hierarchical Bayesian inference of population- and individual-level parameters. **A.** Illustration of backward-in-time coalescent events among six mtDNA samples from a single cell, comparing younger and older individuals. Mutations occur along branches at rate *θ*, and their position on the tree determines the resulting heteroplasmy in the sample. For individuals with higher mitochondrial age *W*, mutations are more likely to be observed at higher heteroplasmy. **B.** Hierarchical Bayesian inference is used to estimate both individual- and population-level parameters for mutation and ageing rates.

### S5.3 Fitting to human data

**Fig. S5.3.**
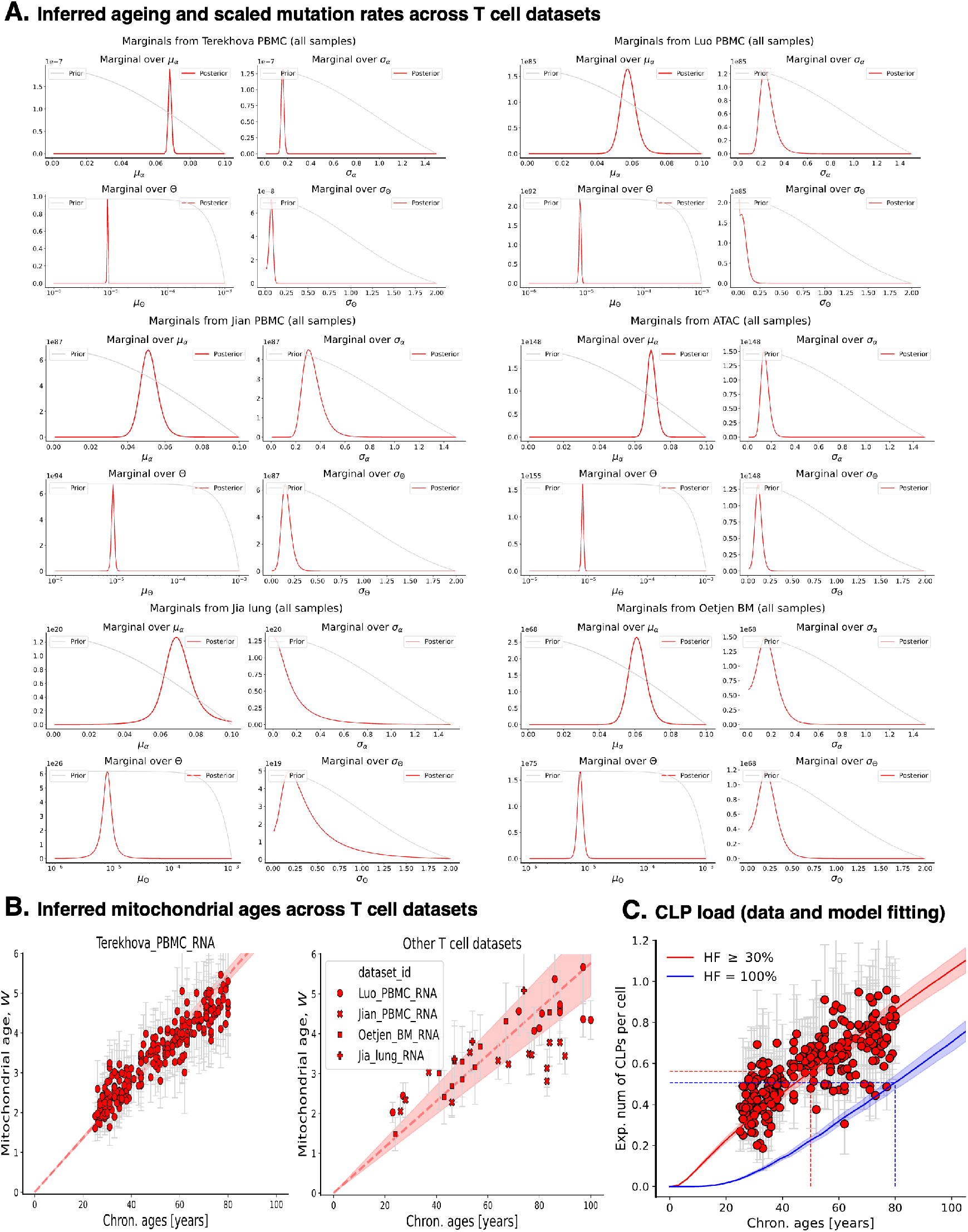
Parameter inference supports appreciable inter-individual variation in mitochondrial ageing and predicts CLP load with age. **A.** Population-level posteriors from the hierarchical model across six T cell datasets. The posteriors for both *µ_α_* and *µ*_Θ_ are broadly consistent across datasets, and almost all datasets show significant non-zero inter-individual dispersion. **B.**Inferred mitochondrial ages for T cells in the Terekhova dataset and the remaining datasets show a clock-like increase with age. Although cell division may contribute to mitochondrial ageing, a simple timescale argument suggests it is insufficient on its own: assuming one division is comparable to one mtDNA turnover generation and mtDNA turnover occurs on the order of one month, even 60 divisions correspond to only ∼5 years of mitochondrial ageing, far less than the age range over which CLP accumulation is observed; **C.** Using the inferred parameters from the Terekhova data, we estimate CLP load over time and compare the model predictions with the observed data (red). We also simulate the trajectory of homoplasmic CLPs (100% heteroplasmy; blue).

### S5.4 Posterior Predictive Checks

**The heteroplasmy distribution of CLP mutations predicted from hierarchical inference matches the observed distribution for each sample.** The following figures illustrate posterior predictive checks to check the validity of our inference model on randomly selected sample IDs from five datasets (for T cells).

**Fig. S5.4.**
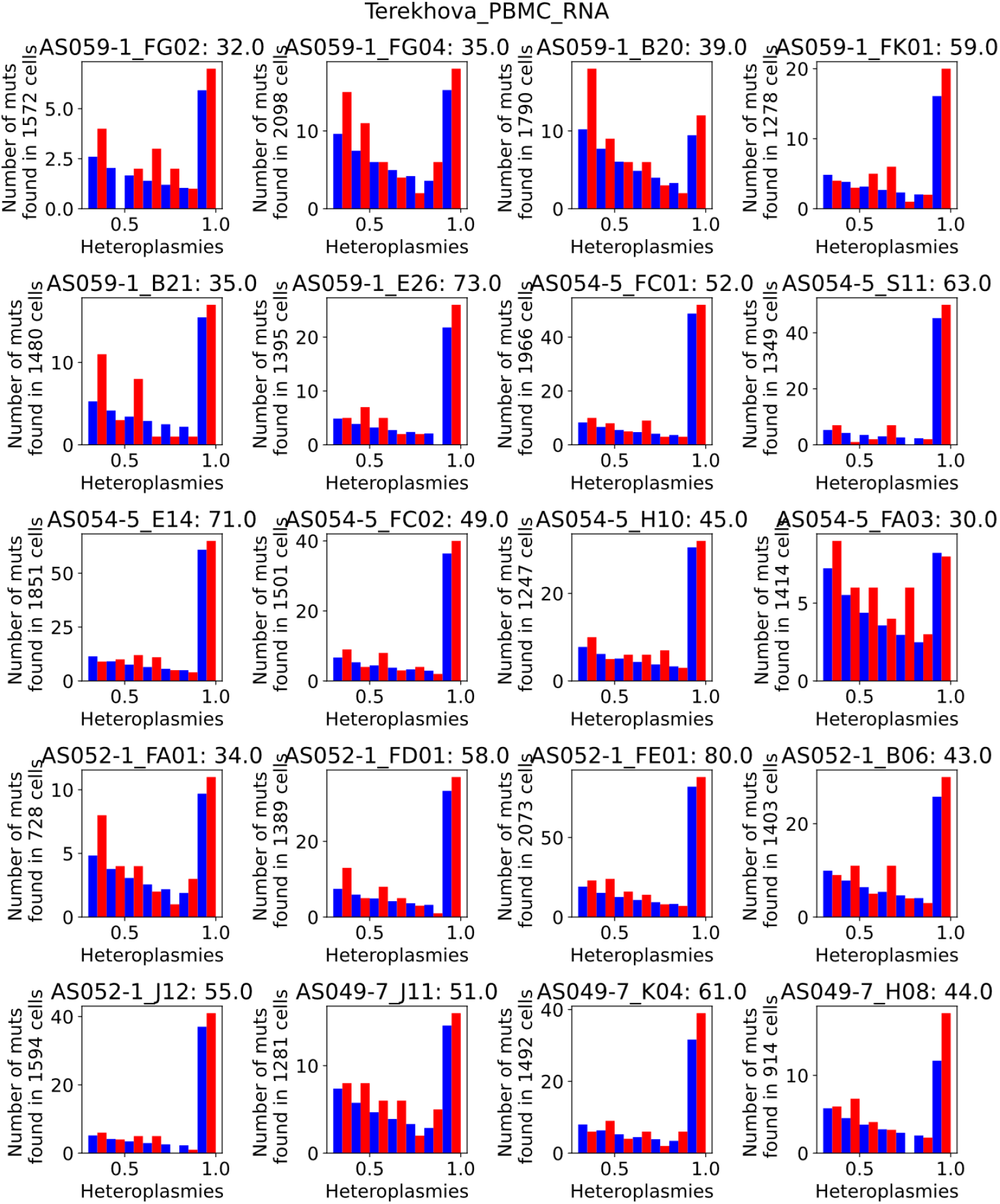
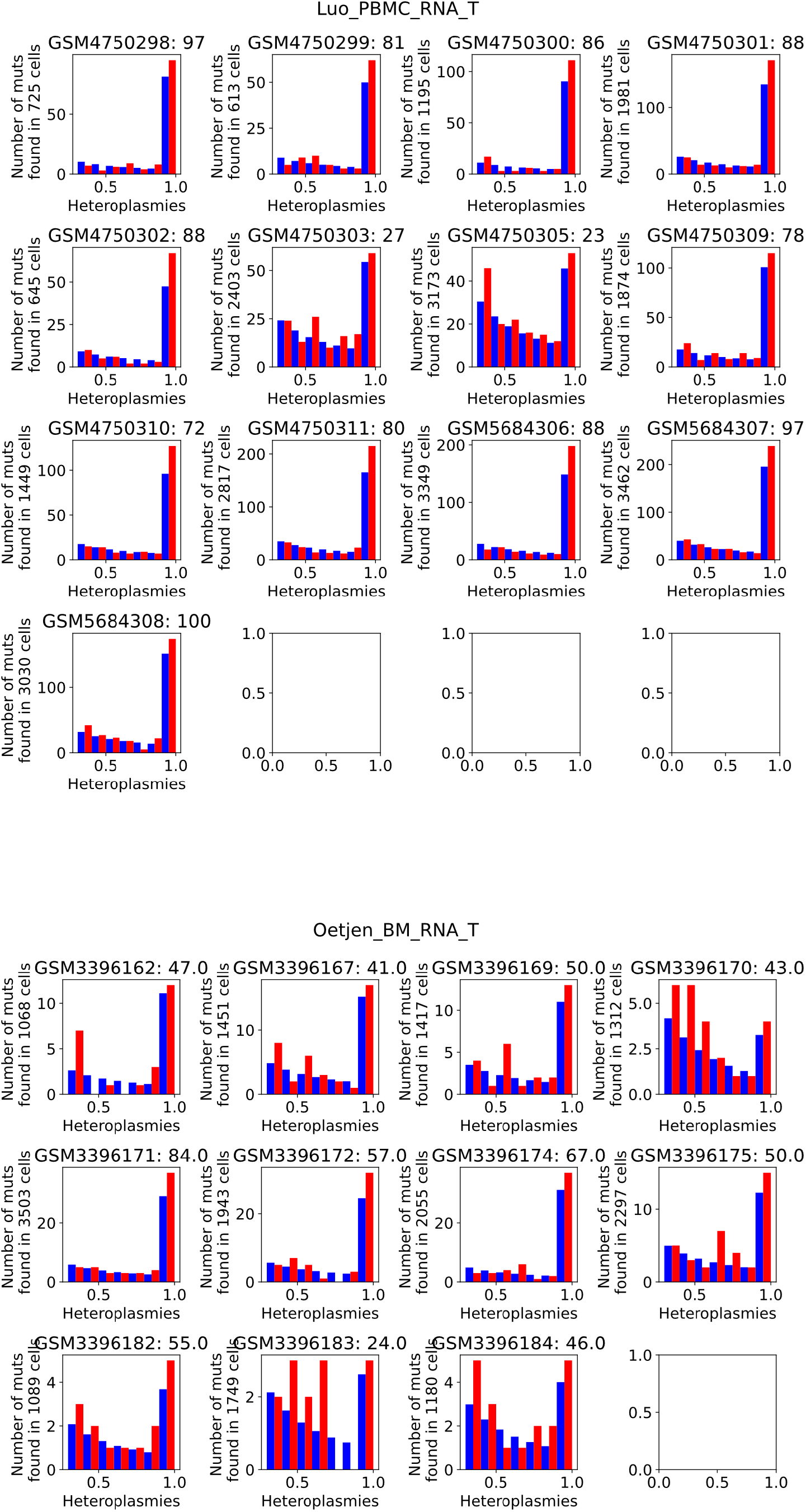
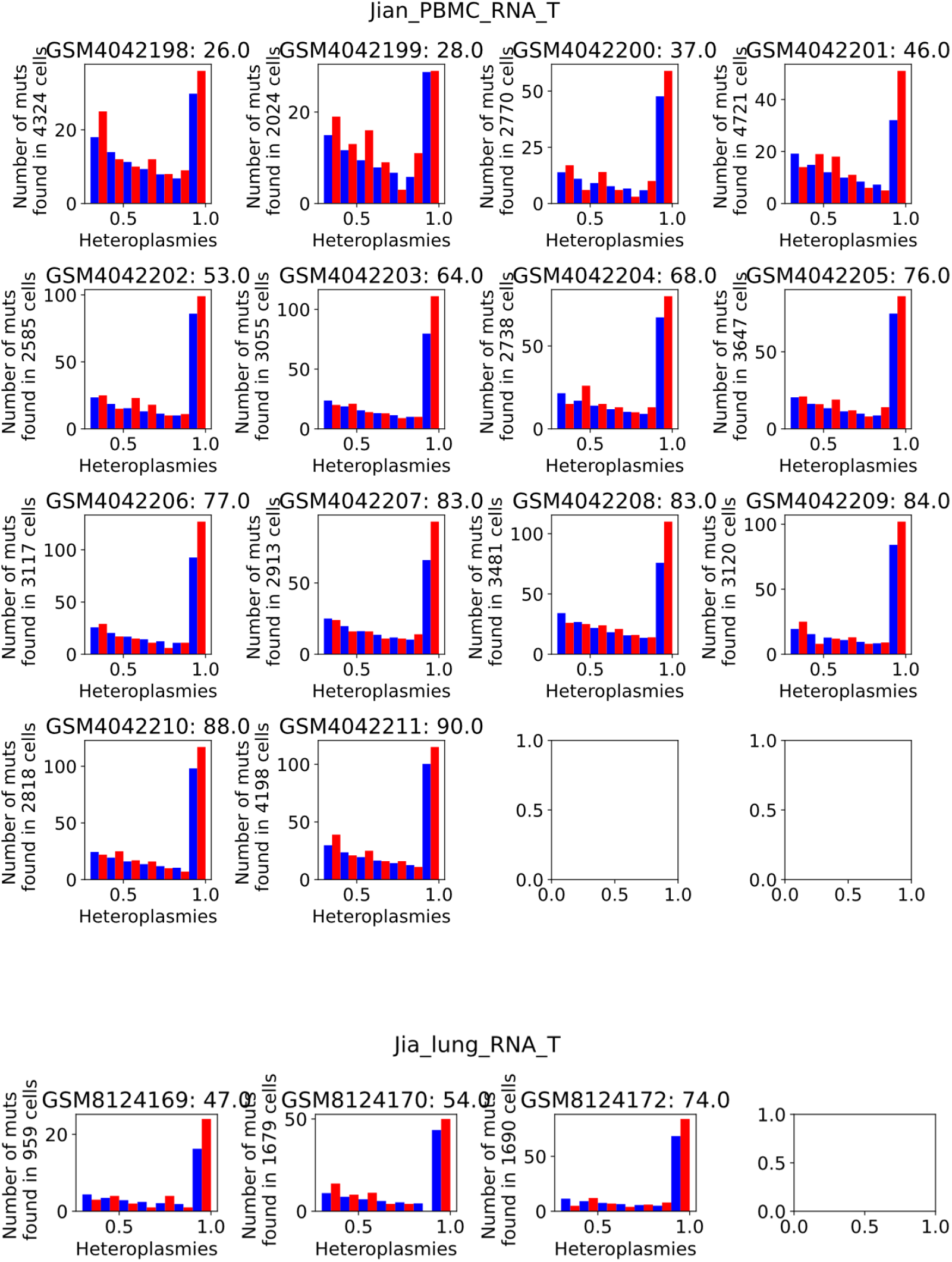
Posterior Checks

## S6 Differential Gene Expression

**Fig. S6.1.**
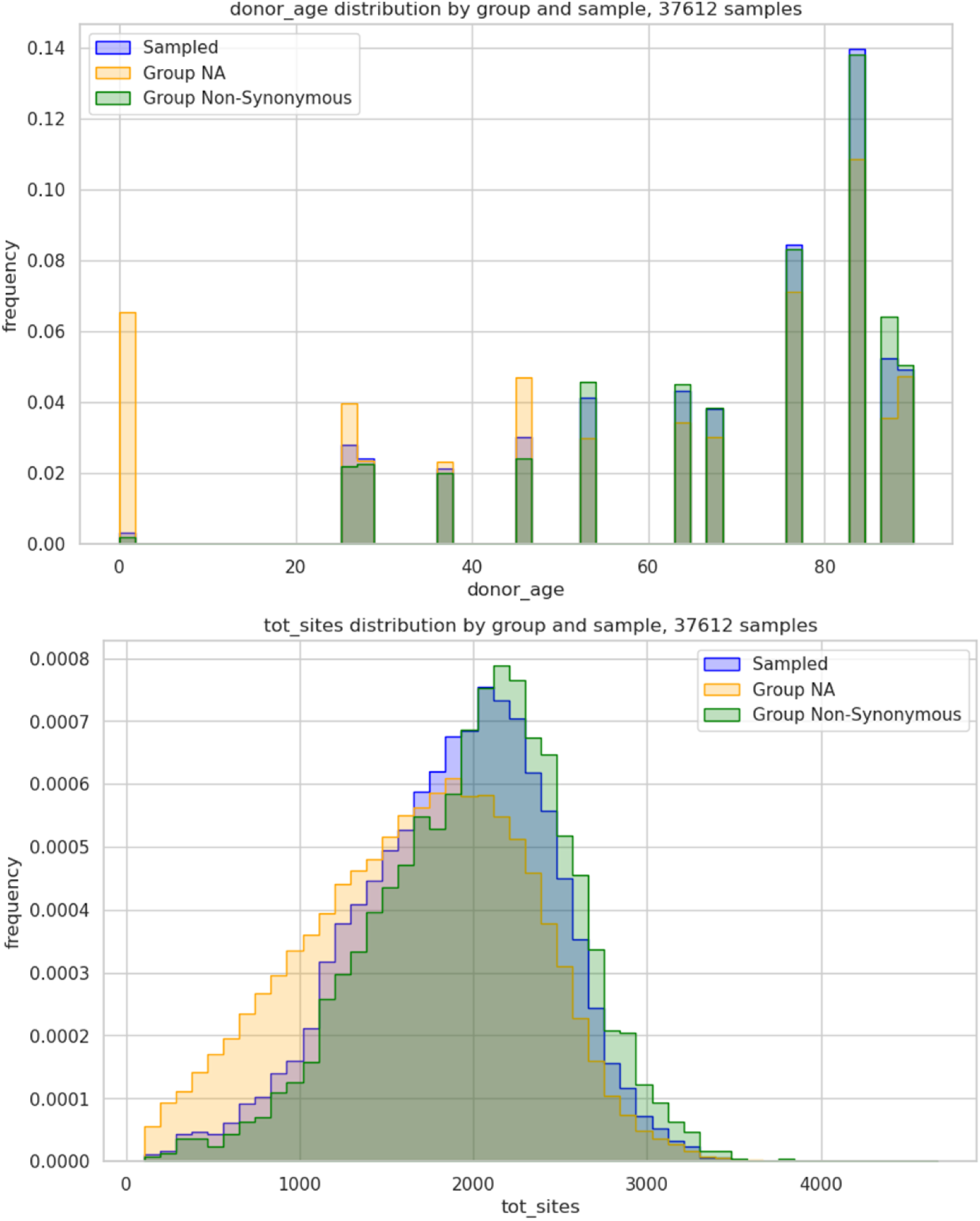
Illustration of the importance-sampling based subsampling procedure showing the marginal distributions of donor age and breadth of coverage after; shown for T-cell data from Jian et al, for a single iteration. We sample from the set of cells with no mutations, or group NA (orange) to match the distribution of cells with at least one Non-Synonymous CLP mutation, or group Non-Synonymous (green). We are interested in matching the marginal distributions of donor age and breadth of coverage, respectively. The sampled set (blue), a subset of group NA, is shown for one random iteration of the sampling procedure. We repeat this procedure 100 times for the results shown in Figure 4.

## S7 Details on sc-ATACseq data

**Fig. S7.1.**
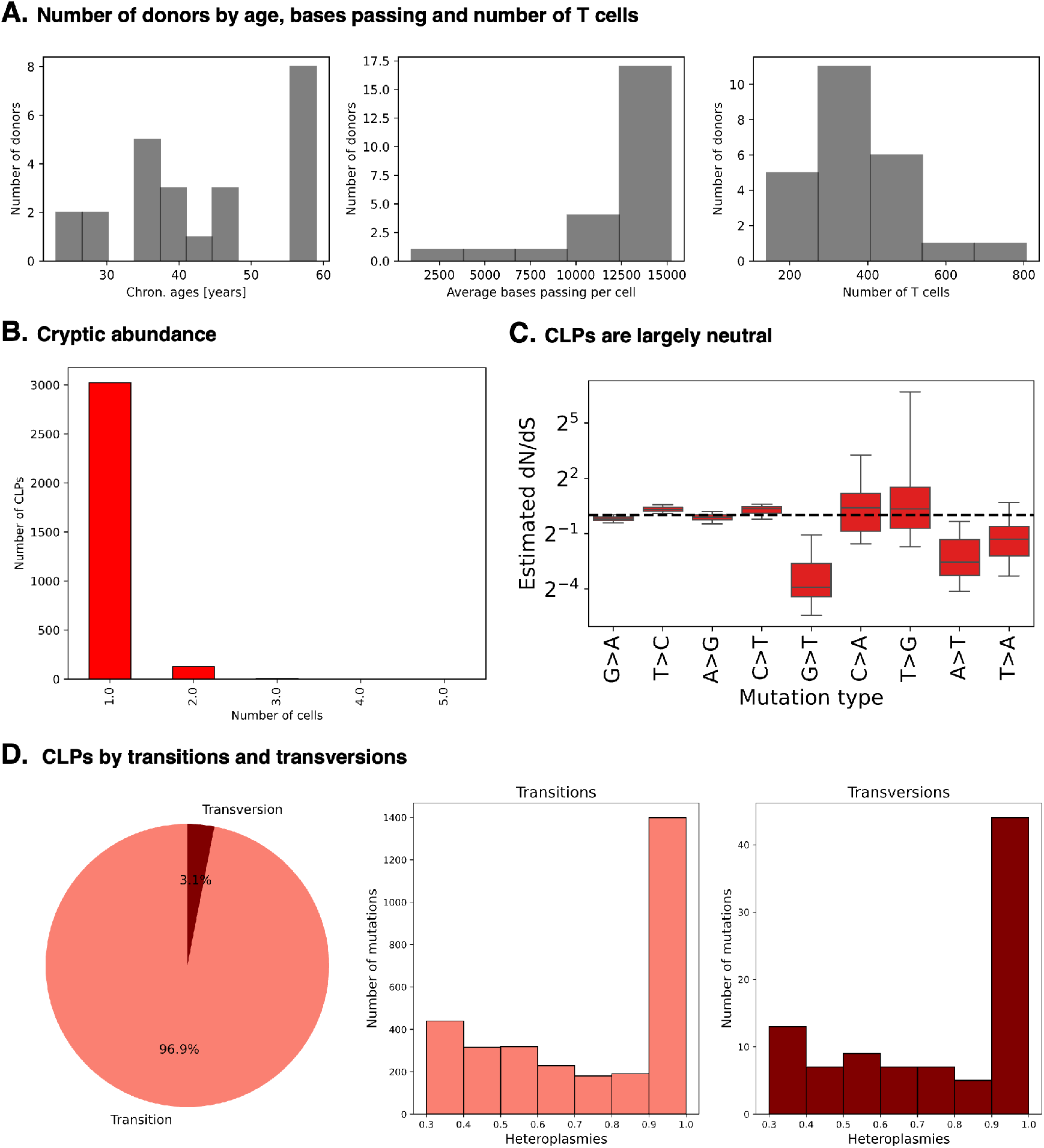
Further corroborations of RNAseq findings by our newly generated mtscATACseq data: **A.** Number of donors by age, bases passing, and number of T cells detected; **B.** Distribution of clone size of CLP mutations, most of which are unique to a single cell; **C.** Inferred dN/dS by type in T cells shows that CLPs are largely neutral; **D.**Most CLPs are transitions, but both are allowed to segregate to higher heteroplasmies, consistent with our RNAseq findings.

---

**Table 1:**
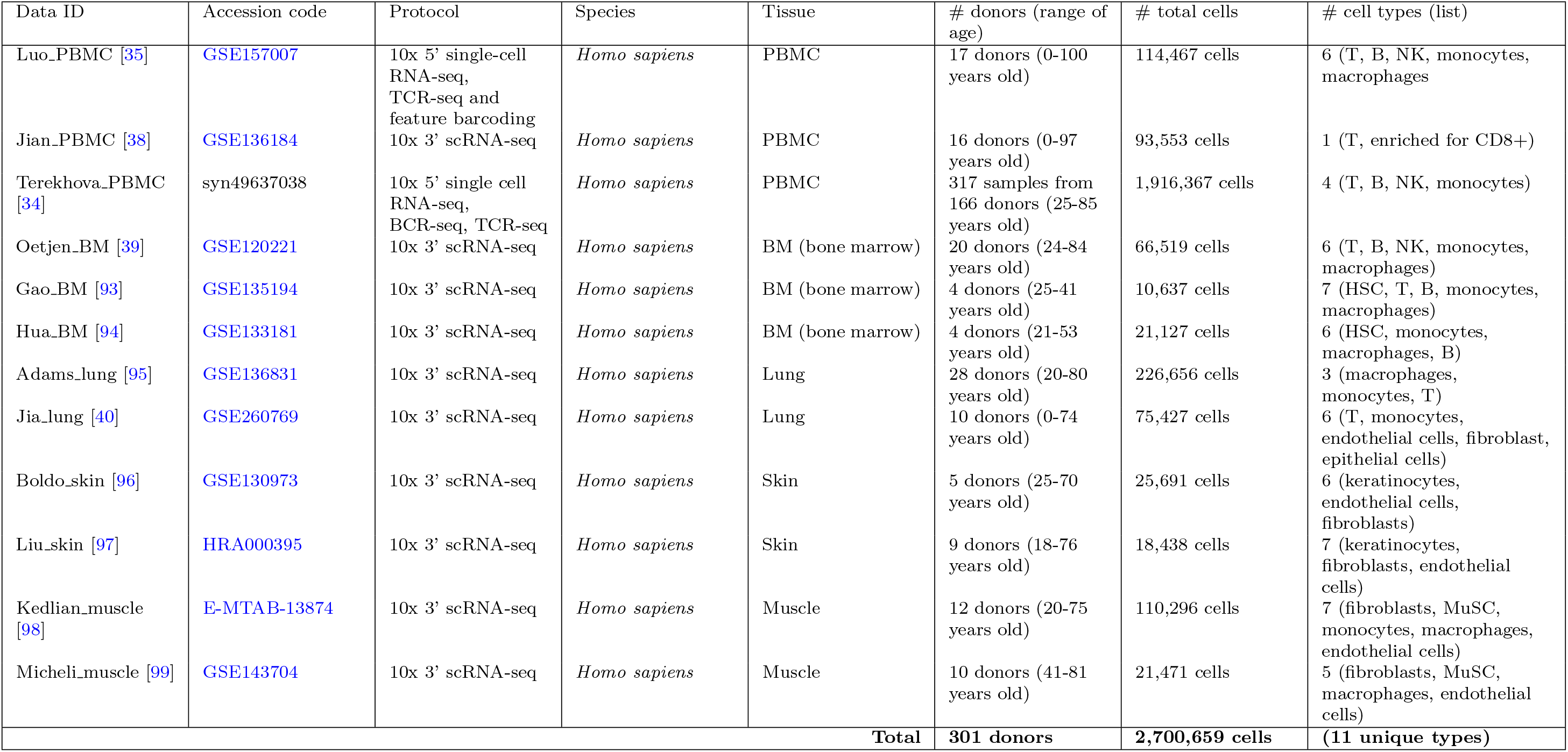
scRNA-seq datasets from human proliferative tissues. The data are generated from 10x Chromium Single-cell Kit 3’ or 5’ protocol, and openly accessible either in GEO, M-TAB, synapse, or HPJA. We also analysed one rats data from [91] with GEO accession number: GSE137869

## Notes

### Competing Interest Statement

The authors have declared no competing interest.

